# Identification of Glucosidase II that regulates STIM1 activation dynamics

**DOI:** 10.1101/2024.04.20.590316

**Authors:** Yangchun Du, Sisi Zheng, Feifan Wang, Jia Li, Xiaoman Yuan, Chao Xi, Yandong Zhou, Youjun Wang

## Abstract

The ubiquitous presence of stromal interaction molecule (STIM), endoplasmic reticulum (ER) Ca^2+^ sensors, plays a crucial role for maintaining Ca^2+^ homeostasis and signaling in mammalian cells by linking ER Ca^2+^ depletion with extracellular Ca^2+^ influx. Although recent advancements have shed light on glycosylation in STIM proteins, further exploration of deglycosylation enzyme modification and its role remains limited by methodological constraints. In this study, leveraging the miniTurbo-driven proximal biotinylation labeling method, we were able to screen for weak and transiently binding protein molecules within specific environments like ER lumina. Herein, we unveil glucosidase II, comprising GANAB and PRKCSH, as a novel partner of the STIM1 complex. Through investigations into Glucosidase II’s (Glu II) regulation of STIM1’s Ca^2+^ affinities both *in cellulo* and *in situ*, alongside its impact on store-operated Ca^2+^ entry (SOCE), we propose a novel mechanism wherein Glu II governs STIM1 activation by modulating Ca^2+^ binding affinity, thereby adjusting the level of activated STIM1 in response to physiological stimulation.

## Introduction

Store-operated Ca^2+^ entry (SOCE) stands as an essential mechanism crucial for both Ca^2+^ signaling and homeostasis ^1–5^. STIM1 and Orai1, residing in the ER and plasma membrane (PM) respectively, spearhead authentic SOCE mediation ^1,6^. Upon ER-Ca^2+^ depletion, STIM1 undergoes a remarkable transformation. The detachment of Ca^2+^ from STIM1’s ER luminal EF-hand motif triggers a conformational shift, unfolding its cytoplasmic domain and releasing the CRAC activation domain (CAD) or STIM1 Orai1-activating region (SOAR) from the Coiled-Coil 1 (CC1) ^7–10^. This event propels STIM1 migration and accumulation at ER-PM junctions, forming characteristic puncta. Here, it engages with and activates Orai channels, precipitating Ca^2+^ influx or SOCE ^1,7,8,11–13^. The implications of STIM1/Orai1 complex malfunctions are profound, implicated in a spectrum of diseases from severe combined immunodeficiency (SCID) to various muscular disorders and inflammatory conditions ^1,14–17^.

Back to 2002, before STIM1 was identified as a key component of SOCE, Williams et al. had confirmed that STIM1 is modified by N-linked glycosylation at two sites within the SAM domain itself, deduced as asparagine residues N131 and N171 ^18^. Despite reports suggesting that STIM1 N-linked glycosylation facilitates EF-SAM destabilization and oligomerization through structural changes in the core α8 helix, in the EF-SAM domain, resulting in enhanced Ca^2+^ influx through CRAC channels ^19^, whether deglycosylation enzyme is involved or further modification of STIM1 glycosylation remains elusive. There remains an important gap in understanding the mechanisms by which Glu II regulates STIM1 activation even after two decades.

While numerous proteins have emerged as regulators of STIM1, influencing its oligomerization, ER-PM translocation, and puncta formation, including notable entities such as POST ^20^, SARAF ^21^, EB1 ^22^, CaM ^23^, ERp57 ^24^, and STIMATE ^25,26^, these regulators typically exhibit high-affinity interactions capable of withstanding harsh cell lysis conditions. Yet, proteins with weak and transient interactions with STIM1 often go unnoticed in conventional techniques. Proximity-dependent biotin identification (BioID) initially unveiled Gelsolin as an interactor of STIM1^27^; however, its extended labeling times (16-18 hours) risk overlooking transient interactions and show diminished efficacy at temperatures below 37°C. In our current investigation, we harnessed the power of miniTurbo enzymes, with TurboID (35 kDa) exhibiting rapid biotinylation within 10 minutes, comparable to BioID’s 18-hour labeling^28–30^. MiniTurbo exhibits maintained specificity and performs optimally at lower temperatures, demonstrating superior activity in the endoplasmic reticulum lumen compared to BioID.

Leveraging advanced proximity labeling techniques (miniTurbo) in tandem with Mass spectrometry (MS), dynamic colocalization assays, and enhanced FRET tools, we unveiled GANAB (the alpha subunit of glucosidase II) and PRKCSH (the beta-subunit of glucosidase II) as novel regulators of STIM1 Ca^2+^-binding affinity. Assessing STIM1’s Ca^2+^ affinity poses formidable challenges. *In vitro* determinations often fall short due to non-physiological conditions, while *in vivo* measurements rely on indirect assessments of STIM1 activity. To surmount these limitations, innovative approaches utilizing molecular tools and biosensors have emerged, enabling real-time monitoring of STIM activities *in situ,* shedding light on its activation mechanisms and Ca^2+^ affinity ^31^. With these cutting-edge tool sets, our current study provides evidence suggesting that Glu II modulates STIM1 activation by influencing its Ca^2+^ affinities, thereby regulating the amplitudes of SOCE. We elucidated the molecular mechanisms underlying Glu II’s regulatory roles. This work not only expands our understanding of STIM1 regulation by deglycosylation via Glu II at the ER side but also holds promise for deciphering broader implications in cellular Ca^2+^ dynamics.

## Results

### The MiniTurbo-driven proximal biotinylation labeling identified Glu II as a partner of the STIM1 interactome

In the process of N-glycosylation, an enzyme known as oligosaccharyltransferase (OST) is recognized for transferring a core structure composed of fourteen sugars (Glc3Man9GlcNAc2) from an ER membrane lipid. Following this initial step, specific enzymes termed glycosidases and glycosyltransferases, operating within the ER, catalyze further alterations, which may involve cleavage or addition of glucose residues ^32^. However, the extent to which glycosidase participates in additional modifications of STIM1 N-glycosylation remains obscure, with little mention of its role in existing literature. Previous research has identified numerous components of the STIM1 interactome ^27^, yet the involvement of any glycosidase remains unverified, likely due to its elusive nature, evading detection under stringent conditions. To address this gap and clarify discrepancies regarding the role of deglycosylation enzymes in STIM1 function, our investigation combines miniTurbo-driven proximal biotinylation labeling with MS analysis. Given that STIM1 exhibits ubiquitous expression across various cell types ^4^, while STIM2 demonstrates heightened expression in brain tissues and dendritic cells ^33,34^, our focus extends to identifying regulators shared between both STIM1 and STIM2. To achieve this, we initially established stable cell lines of HEK293 and Neuro-2A cells expressing miniTurbo-STIM1-YFP and miniTurbo-STIM2-YFP, respectively. Analysis of immunoblots from cell lysates confirmed the successful expression of miniTurbo-STIM1-YFP and miniTurbo-STIM2-YFP (Fig. S1A). Subsequently, the cells were exposed to 50 µM biotin for a duration of three hours to facilitate the biotinylation of the proteome, covering an estimated labeling radius of around 10 nm from STIM1 or STIM2 (Fig. 1A). Following this, the STIM1- or STIM2-proximal proteome was enriched via streptavidin pull-down and resolved using 10% Bis-Tris protein gels. Validation of protein bands was conducted through silver staining and immunoblotting. (Fig. 1B, Fig. S1B). Subsequent MS analysis of three replicated experiments yielded the identification of 140 potential STIM1 interactors and 91 proximal proteins for STIM2, with 22 proteins found to be common to both (Fig. 1C, Fig. S1C-D). Notably, many of these biotinylated proteins were previously known to localize in the endoplasmic reticulum (ER) or on its membrane.

**Fig 1.**
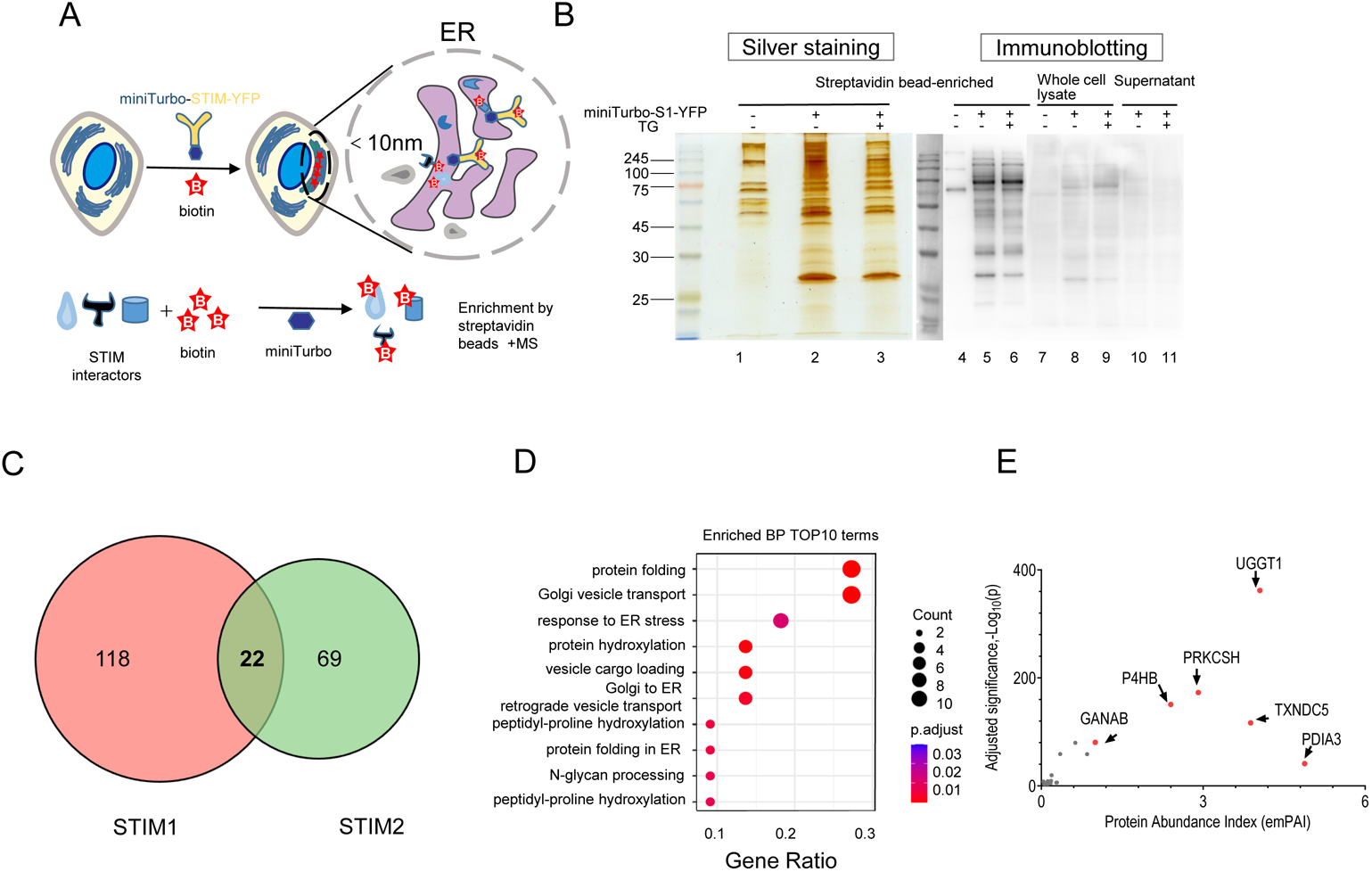
Detection of STIM interactomes utilizing a proximity-labeling approach powered by miniTurbo and mass spectrometry (MS) integration. (A) Illustrated is the workflow for the proteomic mapping strategy aimed at identifying potential STIM interactors within the ER lumen utilizing miniTurbo-mediated biotin labeling *in situ*. Cells expressing MiniTurbo-STIM1/2 were exposed to 50 µM biotin for three hours to facilitate labeling of proteins situated within a 10nm radius of STIM1. Biotinylated proteins were extracted and enriched using streptavidin beads, followed by mass spectrometry (MS) analysis (B). SDS-PAGE revealed silver staining (left) and immunoblotting (right) of biotinylated proteins adjacent to STIM1, with only enriched samples displayed (lanes 1–3). Immunoblots probed with streptavidin-HRP displayed enriched samples (lanes 4–6), whole cell lysates (lanes 7–9), and supernatant samples (lanes 10–11). (C) A Venn diagram is the overlap of proteomics in close proximity to both STIM1 and STIM2. (D) The top 10 enriched biological processes (BP) from proteins proximal to both STIM1 and STIM2 are depicted. (E) A volcano plot showcases STIM-proximal proteomics, with six high emPAI of STIM proteins identified by MS, highlighted as red dots.

We therefore conducted a Gene Ontology (GO) analysis on 118 proteins uniquely associated with STIM1, as depicted in Fig. S1E. This analysis revealed significant enrichment in biological processes such as protein folding and glycosylation. In contrast, the GO analysis for the 69 proteins uniquely linked to STIM2 highlighted significant enrichment in biological processes including actin filament organization, cytoplasmic translation, and actin filament polymerization (Fig. S1F). These findings suggest distinct biological functions for STIM1 and STIM2 in cellular contexts. Further GO analysis of the aforementioned 22 proteins highlighted associations with critical processes like protein folding and N-glycan processing (Fig. 1D). Additionally, a volcano plot illustrating protein abundance and adjusted significance reaffirmed the prominence of six proteins involved in redox reactions (P4HB, TXNDC5, and PDIA3) and N-glycan processing (UGGT1, GANAB, and PRKCSH) (Fig. 1E). For the first time, we provide evidence that the specific glycosidase involved in the STIM1 interactome is glucosidase II, composed of GANAB (alpha subunit) and PRKCSH (beta subunit).

The six STIM interactors we identified can be classified into two enzyme groups engaged in post-translational modifications, one related to redox reactions and the other to N-glycan processing. Considering our initial interests and the documented effects of redox reactions and enzymes such as PDIA3 (also known as ERp57) on STIM1’s Ca^2+^-binding affinity and function ^24^, our current study would still focus on evaluating the influence of glycosylation alterations by Glu II on STIM function, both *in cellulo* and *in situ*.

### The ER luminal domain of STIM1 is a crucial region where PRKCSH or GANAB binds

To further affirm the involvement of Glu II as STIM interactors, we made PRKCSH-mScarlet and GANAB-mScarlet constructs. Our objective is to scrutinize this specific weak interaction using dynamic co-localization techniques with Airyscan super-resolution confocal microscope. Employing two distinct strategies to redistribute STIM proteins, we induced STIM1’s shift from a uniform ER distribution to puncta formation via ER-Ca^2+^ depletion ^4^, and decreased STIM2’s natural puncta formation through intracellular acidification ^35^. Subsequently, we assessed the subcellular localization of PRKCSH and GANAB, both when expressed independently and when co-expressed with STIM. Candidates showing alterations in subcellular distribution only upon co-expression with STIM post-ER Ca^2+^ depletion were identified as dynamic STIM interactors.

First, we test the beta unit of Glu II, PRKCSH. As shown in Fig. 2A, overexpression of PRKCSH alone displays a uniform distribution within the ER during the resting state and maintains a similar distribution following ER depletion induced by ionomycin. However, upon co-expression with STIM1 after ionomycin treatment, PRKCSH undergoes dynamic distributional changes from a uniform ER pattern to punctate structures, as illustrated in Fig. 2B. This synchronized transition, characterized by puncta-like alterations in both PRKCSH and STIM1, signifies an enhanced co-localization (Fig. 2B), supporting the interaction between PRKCSH and STIM1. We further investigated the alpha unit of Glu II, GANAB. Similar to PRKCSH, GANAB also displayed a STIM1-dependent redistribution after ionomycin treatment (Fig. 2C and 2D). The co-localization was significantly enhanced upon ER depletion (Fig. 2D bar chart). STIM2, characterized by a lower affinity for Ca^2+^ compared to STIM1, is adept at detecting minor decreases in ER Ca^2+^ and clustering at the ER-PM junction ^31,36^. Consequently, overexpressed STIM2 displays puncta formation even at rest state, with minimal alteration in distribution upon ER store depletion ^37^. However, acidification induces STIM2 inactivation, resulting in reduced puncta formation ^35^. Consistent with this, GANAB and PKRCSH exhibits a similar STIM2-dependent transition in ER distribution, with its colocalization significantly diminishing upon acidification, as depicted in Fig. S2E&F.

**Fig 2.**
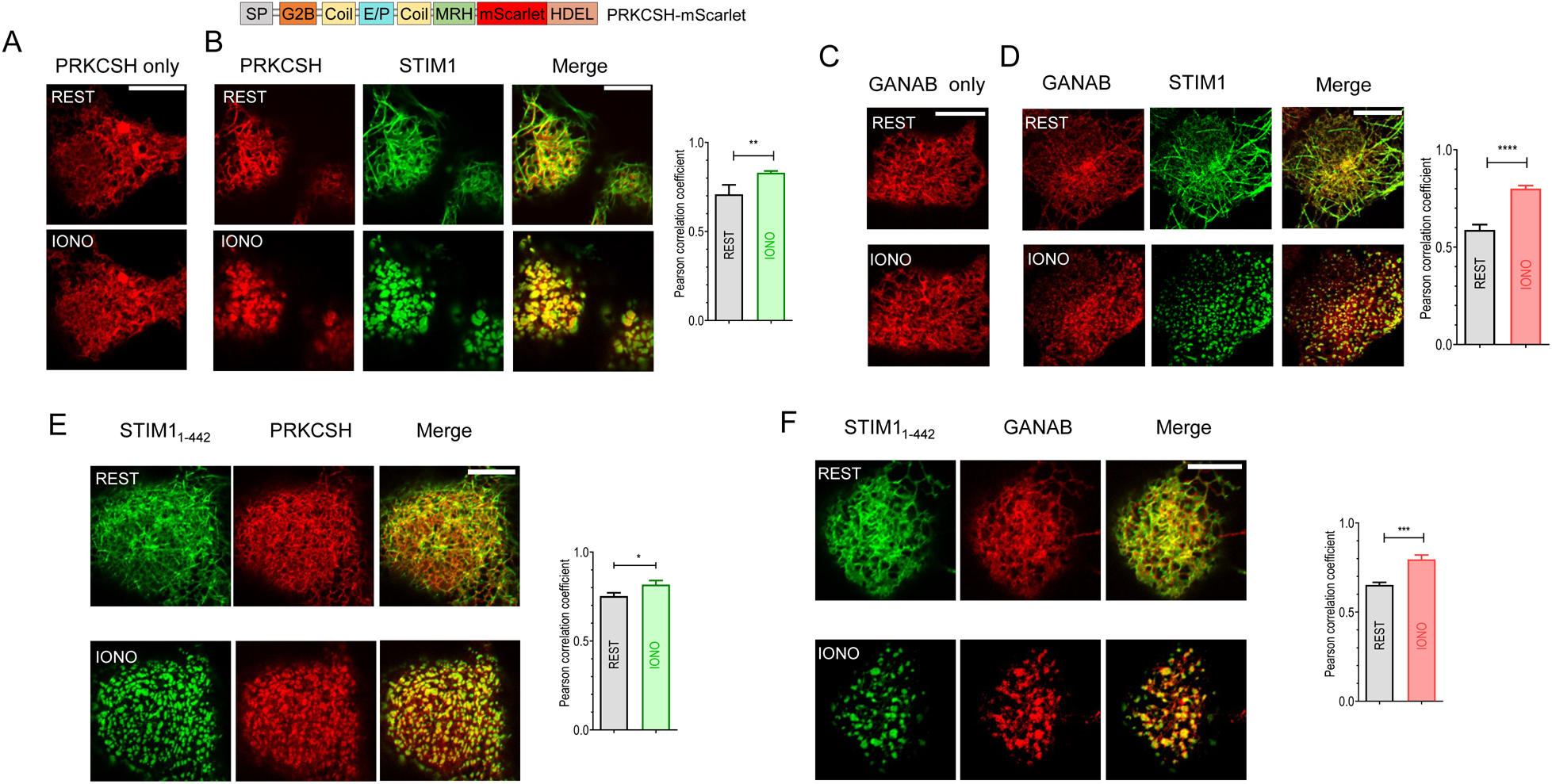
The cytosolic region of STIM1 does not influence the dynamic co-localization of its luminal region with GANAB or PRKCSH. HEK293 cells underwent transient transfection either with GANAB-mScarlet or PRKCSH-mScarlet, either individually or concurrently with wild-type (WT) or truncated variants of mTurquoise2-STIM1. Employing Airyscan super-resolution confocal microscopy, high-resolution images were captured under baseline conditions and after 5 minutes of ER-Ca^2+^ store depletion induced by 2.5 µM ionomycin (IONO). Images were taken from the same field of view. A bar chart on the right of each panel illustrates the calculated Pearson-coefficient, serving as a quantitative measure of co-localization between the specified PRKCSH or GANAB and STIM1 mutants. (A) Sole expression of WT PRKCSH-mScarlet. (B) Co-expression of WT PRKCSH-mScarlet and WT mTurquoise2-STIM1 within the same cells. (C) Sole expression of WT GANAB-mScarlet. (D) Co-expression of WT GANAB-mScarlet and WT mTurquoise2-STIM1 within the same cells. (E) Co-expression of WT PRKCSH-mScarlet and STIM1_1-442_-CFP within the same cells. (F) Co-expression of WT GANAB-mScarlet and STIM1_1-442_-CFP within the same cells. Images are representatives from at least three independent repeats. Scale bar: 10 µm.

As an ER luminal enzyme, Glu II is expected to bind primarily to the luminal portion of STIM1. To investigate this assumption, we generated CFP-tagged STIM1 truncations, including STIM1_1-442_ (lacking the C-terminal cytosolic region) and STIM1_209-685_ (lacking the N-terminal ER luminal region), and evaluated their interaction with mScarlet-tagged PRKCSH using Airyscan super-resolution confocal imaging in HEK293 cells. Remarkably, STIM1_209-685_ exhibited minimal co-localization with PRKCSH (Fig. S2A), while STIM1_1-442_ dynamically co-localized with PRKCSH (Fig. 2E). Additionally, STIM1-C227W ^38^ (a constitutive activation mutation) co-localized with PRKCSH (Fig. S2B).

We conducted similar experiments with GANAB, yielding analogous results. Notably, GANAB exhibited dynamic colocalization with STIM1_1-442_, transitioning from a uniform ER distribution to puncta formation following ER store depletion (Fig. 2F). Conversely, STIM1_209-685_ failed to induce puncta formation with GANAB, even though GANAB already exhibited a punctate distribution at rest (Fig. S2F). Consistently, the constitutive puncta mutation STIM1-C227W, retaining the ER luminal region, successfully induced puncta formation with GANAB (Fig. S2D). These observations align with previous findings, suggesting that the EF-SAM region of STIM1 likely mediates the association with PRKCSH/GANAB, consistent with the localization of STIM1’s glycosylation sites at ASN131 and Asn171^18^.

### The dynamic co-localization of PRKCSH or GANAB with STIM1 relies on the MRH domain of PRKCSH or GANAB-D542

PRKCSH comprises various domains, such as a signal sequence facilitating translocation across the ER membrane, an N-terminal GIIα-binding (G2B) domain, a putative coiled-coil segment, a glutamic acid and proline-rich (E/P) segment, and a C-terminal mannose 6-phosphate receptor homology (MRH) domain, followed by an HDEL signal sequence for ER retention ^39^. Given the essential role of the MRH domain in PRKCSH enzyme activity, our investigation aimed to pinpoint which domain of PRKCSH is crucial for its interaction with STIM1. To address this, we generated mutants of PRKCSH-△MRH-mScarlet (with the MRH domain deleted) and MRH-mScarlet (containing only the MRH domain) and assessed their binding with mTurquoise2-STIM1 and STIM1_1-442_-CFP, respectively. High-resolution images revealed that both mutants were correctly localized in the ER under resting conditions, displaying an ER-like structure (Fig. 3A and 3B, top row images). Following ionomycin treatment, STIM1 puncta failed to recruit PRKCSH-ΔMRH into puncta (Fig. 3A, bottom row), while MRH-mScarlet formed puncta similar to those of STIM1_1-442_ in the same cell (Fig. 3B, bottom row). Taken together, these results suggest that the MRH domain of PRKCSH is required for its association with STIM1.

**Fig 3.**
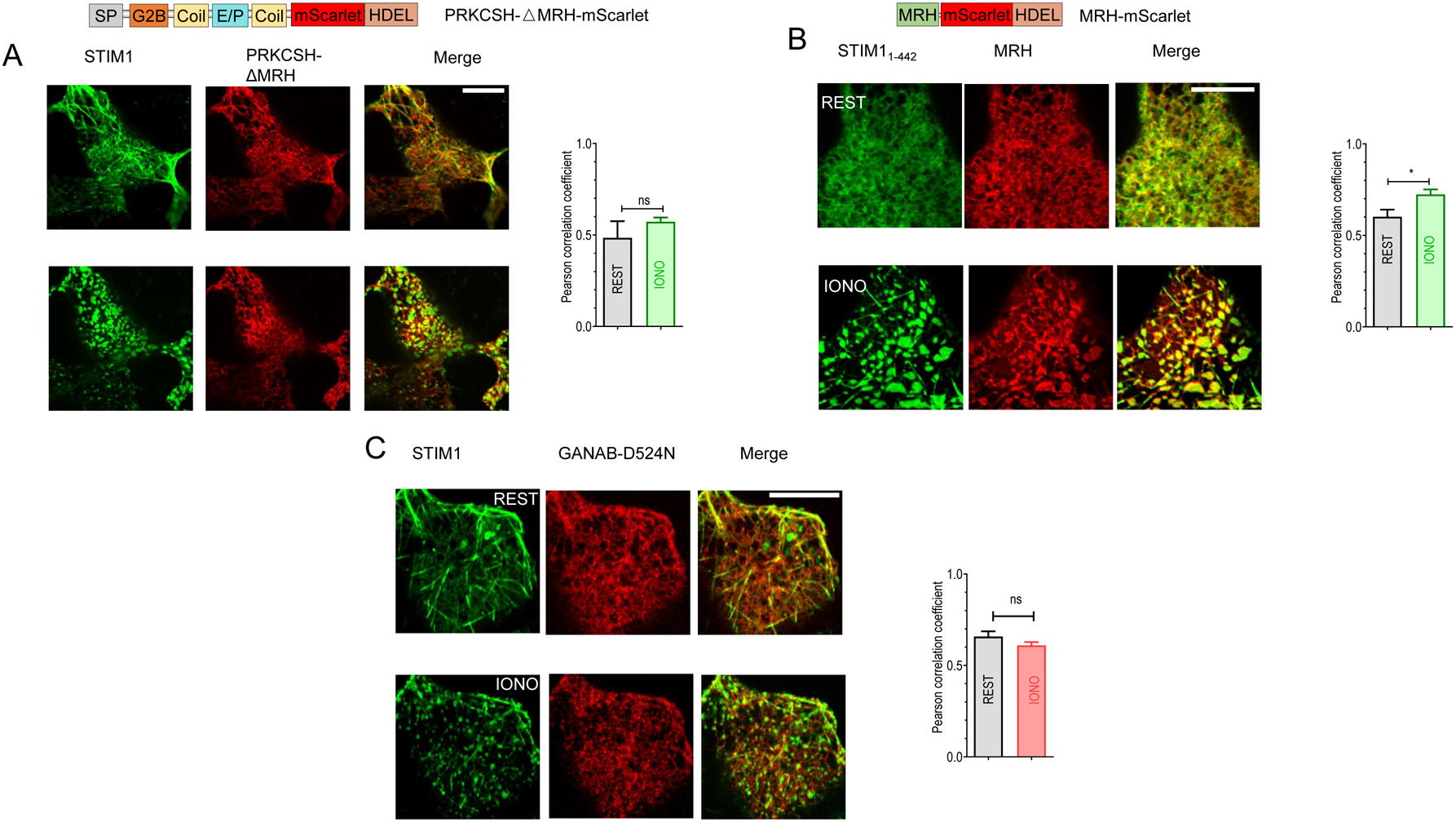
The MRH domain of PRKCSH or the D542 residue of GANAB plays crucial role in their dynamic co-localization with STIM1. HEK293 cells were subjected to transient transfection with various combinations of PRKCSH-△MRH-mScarlet (MRH deletion), or MRH-mScarlet peptide, or GANAB D542N mutation, in conjunction with WT mTurquoise2-STIM1 or STIM1_1-442_-CFP. The bar chart displays the calculated Pearson-coefficient, serving as a quantitative measure of co-localization between the specified PRKCSH or GANAB and STIM1 variants. (A) High-resolution images were obtained depicting the co-expression of PRKCSH-△MRH-mScarlet and WT mTurquoise2-STIM1. (B) High resolution images for co-localization between STIM1_1-442_ and MRH fragment. (C) Examination of colocalization between STIM1 and GANAB-D542N is depicted, with the accompanying bar chart illustrating the statistical analysis revealing the extent of co-localization between STIM1 and GANAB-D542N. Scale bar: 10 µm.

Similarly, in our pursuit to pinpoint the particular domain or amino acids facilitating the interaction between GANAB and STIM1, we focused on the aspartate residue at position 542 of GANAB. Previous research has highlighted the significance of D542 as an essential residue for GluII activity ^40^. We introduced a point mutation, GANAB-D542N-mScarlet, and examined its dynamic co-localization with mTurquoise2-tagged STIM1. Our data demonstrate that while STIM1 effectively transitioned from a uniform ER distribution to puncta distribution upon ER depletion (Fig. 3C, first column), it failed to recruit the GANAB-D542N mutant into puncta (Fig. 3C, second column). These findings emphasize the critical role of the D542 residue of GANAB in modulating its interaction with STIM1.

### *In situ* assessments revealed that non-glycosylable mutation (N131Q-N171Q, 2NQ) in STIM1 increases its Ca^2+^ affinity

We have verified that both components of Glu II undergo the same ER distribution transition alongside STIM1. Within the ER lumen, the EF-SAM domain of STIM1 emerges as a pivotal domain, crucial for Ca^2+^ binding and initiating the first step of STIM1 activation. Previous research has suggested that N-glycosylation of STIM1 impacts the stability of its EF-SAM structure, thereby influencing Ca^2+^ affinity fluctuations ^19^. We posited that further modifications to N-glycosylation by Glu II might similarly affect Ca^2+^ affinity. To assess this, we utilized a FRET assay developed by our team to measure the *in cellulo* Ca^2+^ affinities of STIM1 ^31^. This assay employed engineered STIM constructs localized to the plasma membrane for precise *in cellulo* Ca^2+^ affinity measurements. To circumvent potential artifacts from endogenous STIM1 or STIM2, FRET experiments were conducted in STIM1 and STIM2 double knockout HeLa cells (HeLa SK cells). Our aim was to validate previous *in vitro* findings indicating that N-glycosylation reduces the Ca^2+^-binding affinity of the STIM1-EF-SAM fragment. We compared the *in cellulo* Ca^2+^ affinities of both STIM1 and STIM2 constructs, along with their corresponding 2NQ mutants, using our FRET assay. Unfortunately, the PM-SC1111-2NQ mutation hindered its PM localization, precluding further *in cellulo* evaluation. This observation aligns with previous reports indicating that the N-linked glycosylation sites are crucial for the constitutive cell surface expression of STIM1 ^41^. Notably, the non-glycosylable STIM2 mutation (PM-SC2222-N222Q) did not alter its Ca^2+^ affinity, suggesting minimal impact of N-glycosylation on STIM2’s Ca^2+^ binding (Fig. S3A). This discrepancy may stem from the fact that STIM2 possesses only one glycosylation site at Asn222, unlike STIM1, which boasts two glycosylation sites at Asn131 and Asn171. Consequently, our focus shifted to *in situ* characterization of the Ca^2+^ binding behavior of STIM1 and its non-glycosylable 2NQ mutants.

Direct methods presently available for measuring STIM1’s Ca^2+^ affinities *in situ*, such as CFP-YFP-based FRET pairs employing STIM11-130 and SOAR/CAD/OASF ^9,42^,, or the formation of STIM1 puncta ^43,44^, demonstrate relatively low signal-to-baseline ratios (SBR).This limitation makes it challenging to detect minitor in STIM1’s Ca^2+^ binding behavior. To address this challenge, we developed an improved FRET tool with significantly larger dynamics. In this new STIM1-activation-reporting tool, the fluorescence proteins in STIM1_1-310_-CFP and YFP-SOAR1L were replaced by ECFP△C11 and mNeonGreen△N5 (mNG△N5), respectively. This substitution resulted in a notably expanded dynamic range (1.95 ± 0.15 for CFP-YFP, and 4.89 ± 0.15 for ECFP△C11-mNG△N5), and much higher basal signal (0.23 ± 0.02 for CFP-YFP, and 2.5 ± 0.2 for ECFPΔC11-mNGΔN5). We also avoided indirect measurements with Ca^2+^ sensors by precisely adjusting ER Ca^2+^ levels in permeabilized cells to measure the resulting FRET responses of the new tool (Fig. 4A). Our approach, offering a significantly better signal-to-baseline ratio (SBR) than that achieved with CEPIA1er and STIM1 puncta ^37,43^, provides an accurate assessment of STIM1’s Ca^2+^ affinities. The results showed that the *in situ* Ca^2+^ affinity of STIM1 is 0.69 ± 0.02 mM, with a Hill number of 3.0 ± 0.1 (Fig. 4A). Although this apparent K_d_ value of STIM1 appears higher than in previous reports ^8,43,45,46^, it aligns well with the more accurate measurements of the basal ER Ca^2+^ level (1.3 mM) obtained with our newly developed highly sensitive ER Ca^2+^ indicator, NEMOer ^47^. Thus, based on calculations with the Hill equation, most (87%) STIM1 molecules are bound with Ca^2+^ and remain inactive. The limited fraction (13%) of “activated” Ca^2+^-unbound STIM1 proteins are likely held inactive either by the dominant-negative effect of Orai1 ^48^, or the inhibitory influence of native STIM2 ^49–51^. These inhibitory effects become ineffective in STIM1-overexpressing cells, leading to a subset of cells with constitutive Ca^2+^ entry. This phenomenon, long known but overlooked, is now explained. Overall, our *in situ* results validate the robustness of the STIM1_1-310_ and SOAR1L FRET signals as readouts for STIM activation.

**Fig 4.**
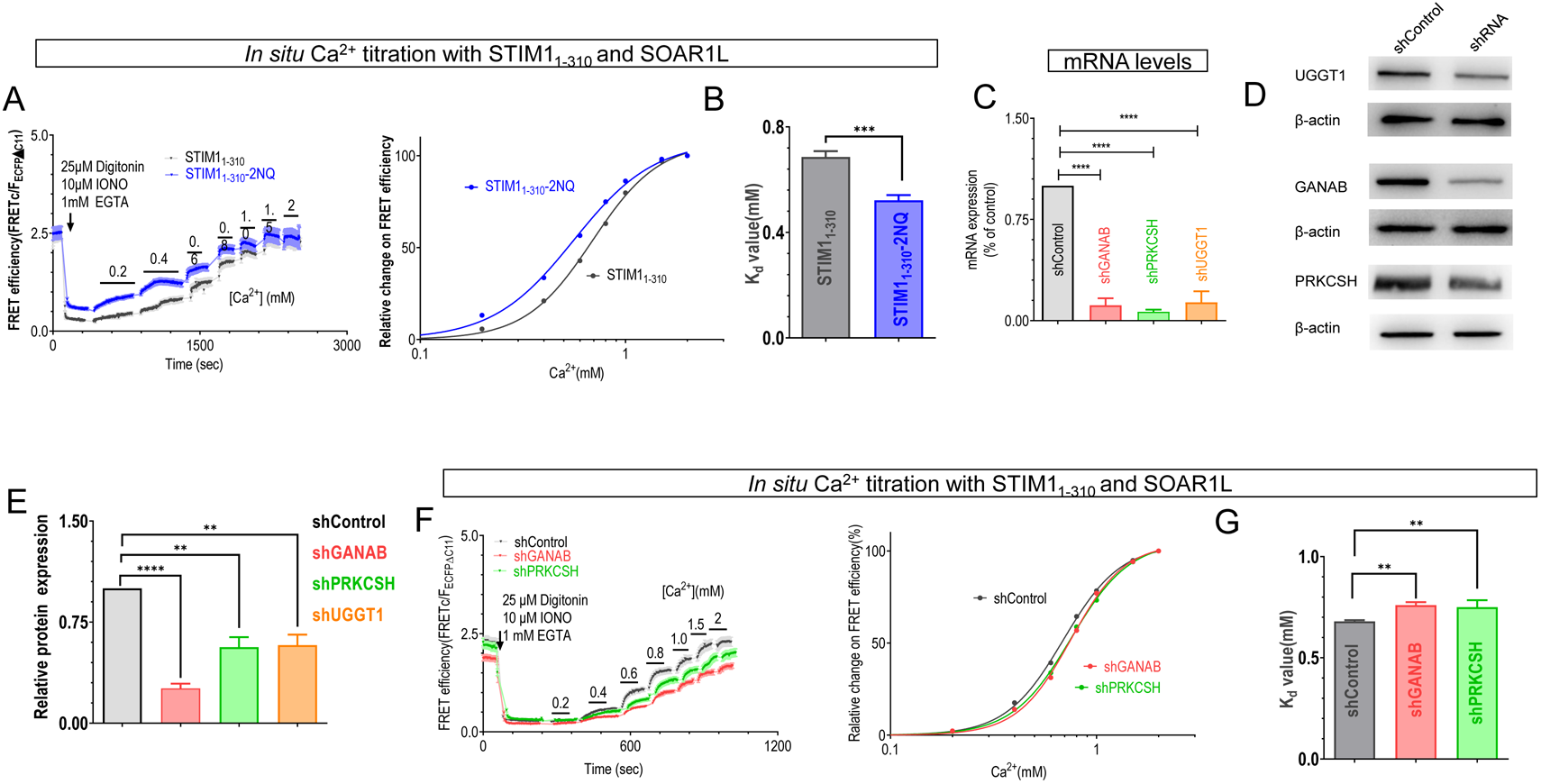
The Ca^2+^ affinity of STIM1 underwent significant alteration upon mutation of its N-glycosylation sites or upon knocking down the expression of *GluII*, which interacts with STIM. To circumvent potential interference from endogenous STIM molecules, HeLa cells with STIM1-STIM2 double knockout (HeLa SK) were employed to express STIM constructs and for subsequent imaging. (A) FRET responses between transiently co-expressed mNG△N5-SOAR1L and STIM1_1-310_-CFP△C11. Left: representative traces; right: Ca^2+^-titration dose response curves (B) Statistical analysis for the Ca^2+^ affinities from data presented in (A) (***, P < 0.0004; paired Student’s t-test) (n = 6, with over 30 cells examined per measurement). (C) Relative mRNA levels of PRKCSH, GANAB, or UGGT1 in cells transfected with non-targeting control shRNA (shControl), shPRKCSH, shGANAB, or shUGGT1 (n = 3, ****, P < 0.0001, unpaired Student’s t-test). (D) Immunoblot analysis of PRKCSH, GANAB, or UGGT1 protein levels in cells transfected with shControl, shPRKCSH, shGANAB, or shUGGT1, and β-actin was used as loading control. (E) Quantification of the identified proteins was conducted for the Western blotting shown in (D). The band intensity of each protein was plotted after normalization to the β-actin signal from the corresponding lane with Image J. (P < 0.003; ****, P < 0.0001, paired Student’s t-test, n = 3). (F) Ca^2+^-titration curves for FRET responses between mNG△N5-SOAR1L and STIM1_1-310_-CFP△C11 constructs in cells transfected with control shRNA, shPRKCSH, or shGANAB. Typical traces are shown in (F) (left, representative traces for FRET response to Ca^2+^ changes; right, Ca^2+^-titration dose response curves). (G) Statistical analysis for the Ca^2+^ affinities from data presented in (F) (**, P < 0.007; unpaired Student’s t-test) (n = 3, with more than 46 cells examined per measurement).

Following this, we evaluated the *in situ* Ca^2+^ affinity of its non-glycosylable mutants in HeLa-SK cells. Initially, we investigated the impact of the N131C-N171C (2NC) mutation. STIM11-310-2NC exhibited a very low basal FRET signal with SORAL and minimal response to changes in Ca^2+^ concentration, indicating a low Ca^2+^ affinity with constitutive activity (Fig. S3C, S3D, and S3E). This is consistent with a previous *in vitro* report, which showed that the STIM1-EF-SAM fragment with the 2NC mutation had a K_d_ value around 4.4 mM ^19^. While these findings underscore the importance of glycosylation sites, we attribute this significant change in Ca^2+^ affinity to possible redox responses of the two introduced cysteine residues. Subsequently, we examined the Ca^2+^ binding behavior of STIM11-310-2NQ. The measured K_d_ value (0.55 ± 0.01 mM) for this non-glycosylable mutant, with no extra cysteine residues, was significantly lower than wild-type (WT) STIM1_1-310_ (Fig. 4A and 4B), indicating that completely deglycosylation alone enhances STIM1’s Ca^2+^ affinity.

### The lowering of native deglycosylation enzymes results in a decrease in STIM1’s Ca^2+^ affinity

Glu II is responsible for removing alpha-1-3-linked glucose from high mannose oligosaccharides, rather than completely abolishing glycosylation. Our inquiry centers on whether this N-glycosylation modification still influences STIM1’s Ca^2+^ affinity. Initially, we scrutinized the impact of UGGT1, PRKCSH, or GANAB overexpression on FRET signals between STIM1_1-310_ and SOAR1L, acting as indicators for STIM1 activation. Intriguingly, the overexpression of these proteins did not alter STIM1’s K_d_ value (Fig. S3F), suggesting that endogenous levels of these proteins suffice for N-glycosylation processing activities. Another plausible explanation could be that overexpressing just one unit of Glu II may not sufficiently augment its expression level. However, due to the complexity of transiently co-expressing four constructs (STIM11-310, SOAR1L, GANAB, and PRKCSH), we encountered challenges in successfully obtaining healthy cells expressing all four proteins. Consequently, we proceeded to explore the effects of knocking down these proteins using shRNA technology.

The efficiency of the knockdown was assessed through quantitative RT-PCR, revealing a significant decrease in their mRNA levels (Fig. 4C). This was further confirmed by Western blot analysis, demonstrating a notable reduction in the expression of these proteins (Fig. 4D-E). Interestingly, knocking down UGGT1 did not affect the Kd value of STIM1 (Fig. S3G). This could be due to compensatory mechanisms within the cells or the presence of alternative regulatory processes such as UGGT2 ^32,52^ However, after knocking down GANAB or PRKCSH, the K_d_ value of STIM1 significantly increased (shControl: 0.68 ± 0.02mM; shGANAB: 0.77 ± 0.01mM; shPRKCSH: 0.78 ± 0.01mM) (Fig. 4F-G), suggesting that the knockdown of GANAB or PRKCSH may weaken STIM1’s ability to bind Ca^2+^. To assess the specificity and the role of STIM1’s glycosylation site in this regulation, we examined the impact of knocking down PRKCSH or GANAB on the K_d_ value of STIM1-2NQ, which lacks oligosaccharides. The results indicate that knocking down PRKCSH or GANAB does not alter the K_d_ value of STIM1-2NQ (Fig. S3H). Additionally, the STIM1-2NQ mutation abolishes its ability to recruit PRKCSH or GANAB into puncta, despite still exhibiting the transition from uniform ER distribution to puncta (Fig. S4A and S4B). These findings suggest that GANAB and PRKCSH modulate the Ca²⁺ affinity of STIM1 through alterations in its glycosylation state.

### The knockdown of PRKCSH or GANAB promotes STIM1 activation and SOCE

Upon depletion of ER stores, calcium ions dissociate from the EF hands of STIM1, triggering a structural alteration in the EF-SAM domain located in the ER lumen. This alteration induces a conformational switch in the cytosolic region of STIM1, leading to the detachment of the CAD/SOAR domain from CC1. Consequently, the CAD/SOAR domain projects towards the plasma membrane (PM), where it interacts with and regulates the activity of Orai channels. Building upon these findings, we observed a significant decrease in the Ca^2+^ affinity of STIM1 upon knockdown of PRKCSH or GANAB (Fig. 4F), suggesting a potential impact on its transition from a resting to an activated state. Therefore, we investigated the effect of knockdown on the activation steps of store-operated calcium entry (SOCE). However, neither overexpression nor shRNA knockdown of GANAB, PRKCSH, or UGGT1 significantly affected the FRET signals between CFP-STIM1 and YFP-STIM1 (Fig. S5A and S5B) or between STIM1-YFP and Orai1-CFP at rest (Fig. S5C and S5D). These results indicate that these proteins do not influence STIM1 oligomerization or the coupling of STIM1 with Orai1.

We proceeded to delve deeper into the functional implications of STIM1’s interactions with PRKCSH or GANAB within the cellular milieu. Initially, we examined the effect of GANAB or PRKCSH on STIM1 functionality. HEK293 cells, stably expressing the highly sensitive cytosolic Ca^2+^ indicator TurNm (TurNm cells) ^53^, were transiently transfected with shRNA targeting PRKCSH or GANAB. Following passive depletion of the ER Ca^2+^ store using thapsigargin (TG), which induced a transient release of Ca^2+^ from the ER lumen, 1 mM extracellular Ca^2+^ was added back to allow Ca^2+^ influxes via store-operated calcium entry (SOCE). Compared to blank controls, knockdown of GANAB or PRKCSH significantly augmented SOCE (Fig. 5A). This finding aligns with the increase in the Kd value of STIM1 observed upon knocking down PRKCSH or GANAB (Fig. 4F). Subsequently, we investigated SOCE under sub-maximal stimulation mimicking physiological conditions. We assessed the formation of STIM1 puncta induced by submaximal activation of muscarinic acetylcholinergic receptors with 10 μM carbachol (CCh). Following knockdown of either PRKCSH or GANAB, a notable increase in the number of cells forming STIM1 puncta in response to 10 μM CCh was observed compared to blank controls (Fig. 5B). This observation correlates with the findings regarding SOCE induced by 10 µM CCh (Fig. 5C). Overall, knocking down PRKCSH or GANAB renders STIM1 more readily activated.

**Fig 5.**
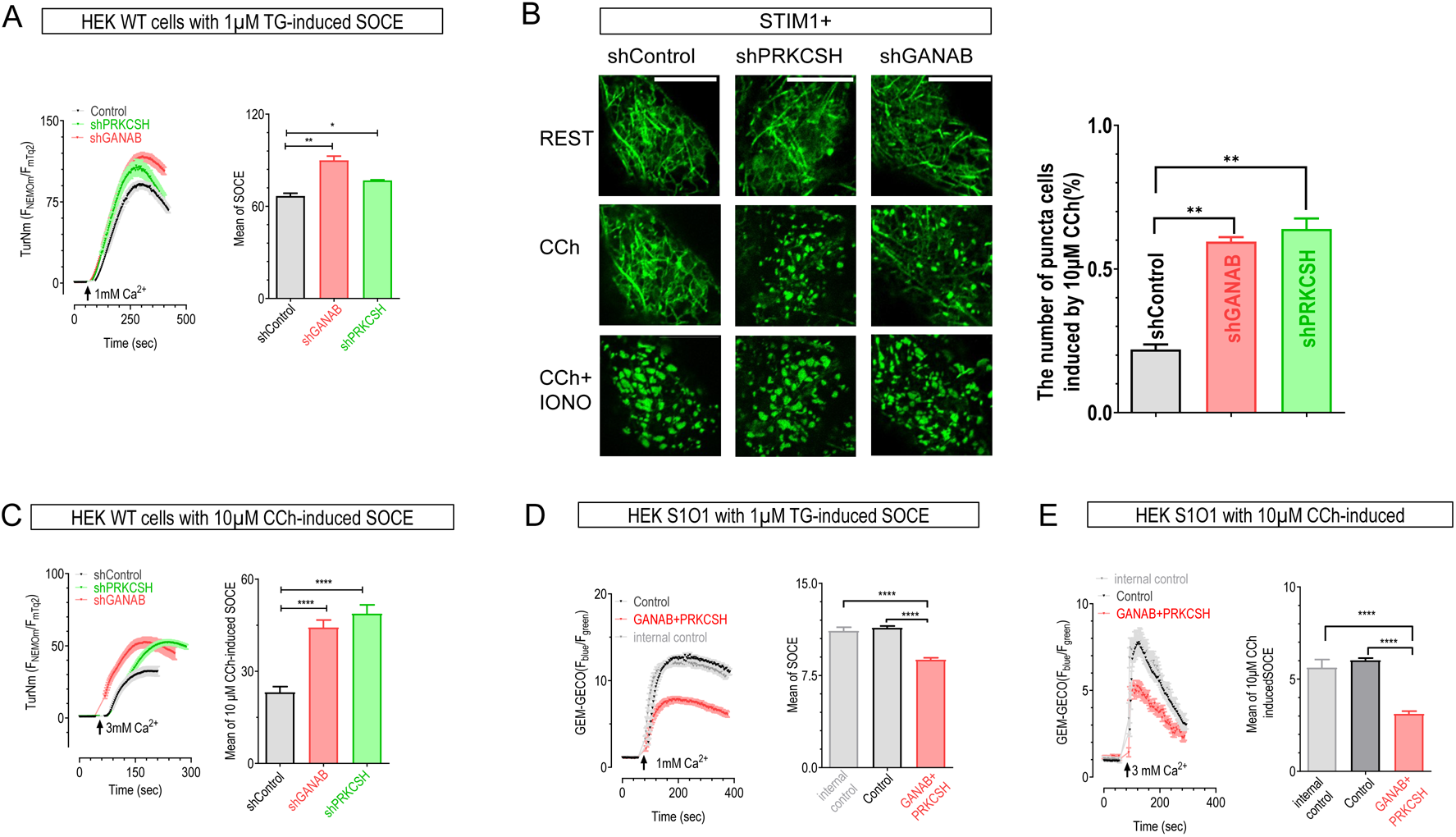
Modulation of TG or CCh-induced SOCE by Glucosidase II. HEK293 cells, stably expressing the highly sensitive cytosolic Ca^2+^ indicator TurNm (TurNm cells), were transiently transfected with different shRNAs or PRKCSH variants. Before recordings, ER Ca^2+^ stores were depleted by incubating cells in nominally Ca^2+^-free solutions with 1 μM TG or 10 μM CCh for 10 minutes as indicated. Left: representative traces; right: statistical analysis. (A) representative trace for Store-operated Ca^2+^ entry (SOCE) responses induced by TG. (n = 4, >50 cells per measurement, **, P < 0.001; *, P < 0.01, paired Student’s t-test). (B) Confocal images showing mTurquoise2-STIM1 distribution in cells co-transfected with mTurquoise2-STIM1 and different shRNAs. Images captured at basal, 5 minutes after 10 µM Carbacholine (CCh) stimulation, and 5 minutes after subsequent addition of 2.5 µM ionomycin (IONO). Right: statistical analysis (n = 3, >20 cells per measurement, **, P < 0.001, Student’s t-test, scale bar, 10 µm). (C) representative trace for SOCE induced by 10 μM CCh. TurNm cells transiently transfected with control shRNA, PRKCSH shRNA and GANAB shRNA, respectively. Cells pre-incubated in nominally Ca^2+^-free solutions with 10 μM CCh for 5 minutes (n = 3, >50 cells per measurement, ****, P < 0.0001, Student’s t-test). (D) representative trace for SOCE induced by 1 μM TG in HEK cell line stably expressing STIM1-YFP and Orai1-CFP, transiently co-expressing WT PRKCSH and WT GANAB. Blank control is no transient expression of exogenous DNA. Internal control is transiently expressing either PRKCSH or GANAB. (n = 3, >30 cells per measurement, ****, P < 0.0001, Student’s t-test). (E) representative trace for SOCE induced by 10 μM CCh in HEK cell line stably expressing STIM1-YFP and Orai1-CFP, transiently co-expressing WT PRKCSH and WT GANAB. Blank control is no transient expression of exogenous DNA. Internal control is transiently expressing either PRKCSH or GANAB. (n = 3, >30 cells per measurement, ****, P < 0.0001, Student’s t-test).

Furthermore, we aim to investigate whether the overexpression of PRKCSH and GANAB yields effects contrary to knockdown strategies, which typically inhibit store-operated calcium entry (SOCE) in HEK cells. As previously discussed, the presumption of sufficient endogenous PRKCSH and GANAB levels in HEK cells rendered us unable to discern any alterations in Ca^2+^ affinity within STIM1 through the overexpression of either PRKCSH or GANAB via *in situ* assessments. To counteract the potential influence of endogenous PRKCSH and GANAB on the limited native levels of STIM1, we conducted our experiments on HEK cells (HEK S1O1 cells) stably expressing exogenous STIM1 and Orai1. This approach ensures an abundance of STIM1 within the endoplasmic reticulum (ER) membrane and an excess of Orai1 channels in the plasma membrane.

GANAB WT and PRKCSH WT were co-transfected into HEK S1O1 cells, followed by the Ca^2+^ measurement and comparison of SOCE among these groups. As depicted in Figure 5D, in PRKCSH/GANAB group, the assembly of normal functional Glu II significantly inhibited SOCE induced by TG compared to the blank control. Within our experimental framework, the internal control comprises cells expressing solely either GANAB or PRKCSH within the same field of measurement. As depicted in Figure 5D, the individual expression of either GANAB or PRKCSH alone failed to manifest any inhibitory effect on store-operated calcium entry (SOCE), similar as blank control. In this context, the blank control mirrors the effect of knocking down the entire Glu II compared to the overexpression of GANAB WT and PRKCSH WT. With limited endogenous Glu II, the N-glycosylation of the overexpressed STIM1 remains unaltered by Glu II. Consequently, this results in more STIM1 remaining in an activated state under similar stimulation, leading to the activation of more Orai1 channels on the plasma membrane and enhancing SOCE. We were also keen to explore whether the inhibition persists under conditions of submaximal ER store depletion induced by 10 µM CCh. And co-expression of PRKCSH and GANAB indeed demonstrated a notable inhibition of SOCE compared to both the blank control and the internal control, as illustrated in Figure 5E. In concert, the plentiful presence of PRKCSH and GANAB collaboratively regulates the dynamics of STIM1 activation in response to stimuli by mediating its calcium affinity (Fig. 6).

**Fig 6.**
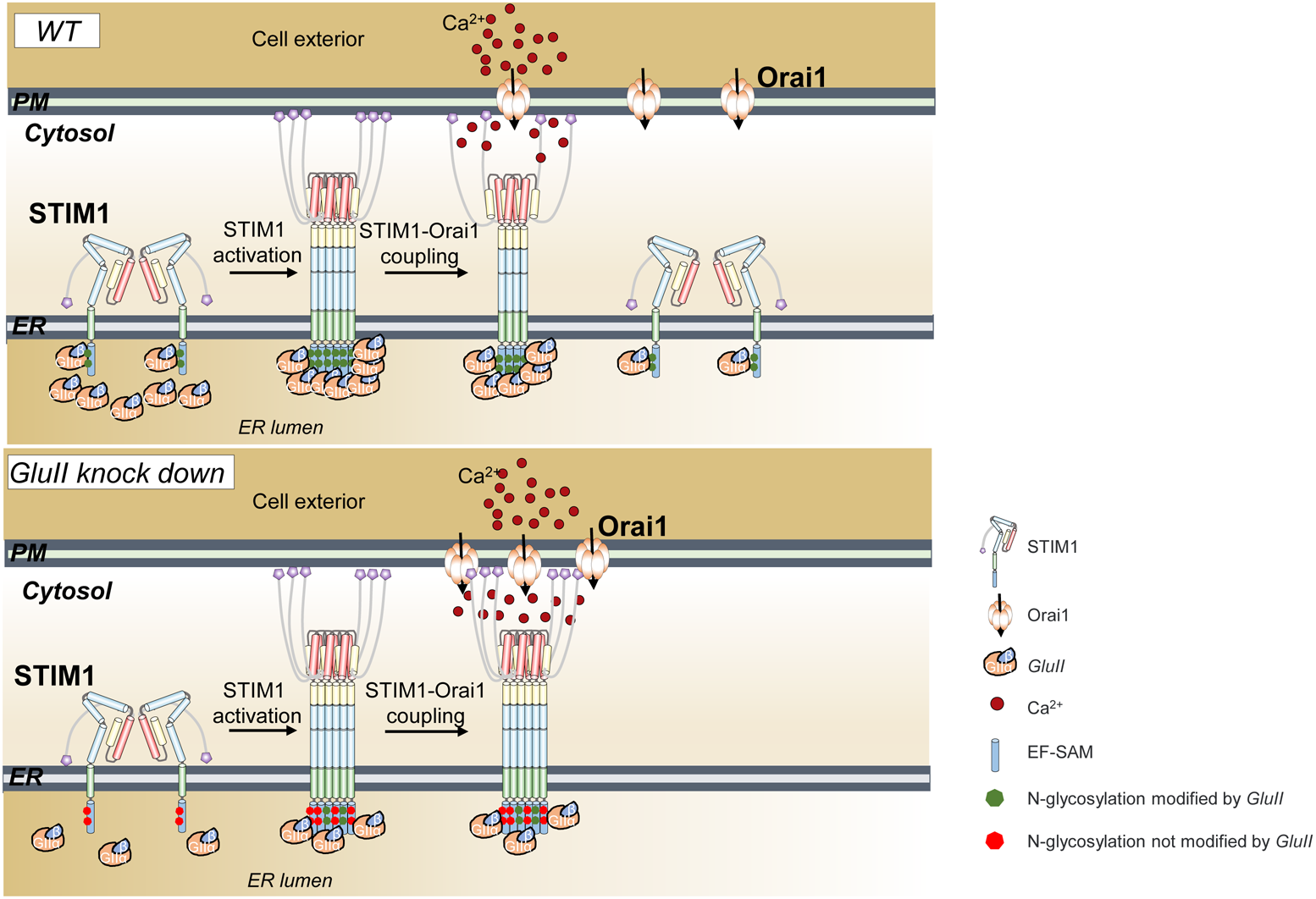
Model for the regulation of STIM1 activation by Glucosidase II and its physiological role in the generation of Ca^2+^ signals. Top: Under physiological conditions, abundant Glu II modifies all STIM1 proteins at their N-glycosylation sites. Store depletion induces the activation of STIM1 proteins, with further co-localization of Glu II with STIM1 securing these modifications. This process significantly increases its Ca^2+^ binding affinity, resulting in less STIM1 remaining in an activated state. By enhancing the Ca^2+^ binding affinity of STIM1, Glu II regulates SOCE to prevent Ca^2+^ overload. Bottom: Under pathological conditions, represented here by Glu II knockdown, the N-glycosylation sites of STIM1 are not fully modified due to insufficient Glu II levels. This leads to more STIM1 activation under a similar decrease in ER Ca^2+^ which results in enhanced SOCE as shown in Fig 4B. Our study indicates that Glu II intricately adjusts STIM1’s response to ER Ca^2+^ variations through targeted modifications at its N-glycosylation sites, influencing its activation status.

## Discussion

Calcium signaling stands as a pivotal regulator of numerous cellular functions, meticulously governed by an array of pathways, channels, and calcium pumps. Among these, store-operated calcium entry (SOCE) emerges as a cornerstone pathway, orchestrated by the interplay of STIM and Orai proteins^1^. SOCE represents a ubiquitous signaling event observed in both excitable and non-excitable cells, serving as a linchpin in a myriad of physiological and pathophysiological responses ^54^. Extensive studies have elucidated the multifaceted regulation of STIM through various post-translational modifications (PTMs), encompassing phosphorylation ^55^, acylation ^56,57^, glycosylation ^18,19^, and oxidation by reactive oxygen species (ROS) ^58,59^. Although the N-glycosylation of STIM1 was first reported in 2002^18^ there has been a notable lack of further investigations into the potential involvement of deglycosylation enzymes in modulating STIM1 glycosylation over the past two decades. Additionally, the specific enzyme(s) responsible for this process remain unidentified.

One potential explanation lies in methodological limitations. Following the identification of STIM1 as the Ca^2+^ sensor ER for SOCE^60–62^, numerous STIM1 interactors were initially identified through conventional biochemical methods^24,63,64^, which are capable of detecting only robust interactions resilient to harsh lysis conditions. The advent of enzymatic proximity labeling has revolutionized the study of local protein ensembles^65,66^, enabling the discovery of protein-protein interactions, whether they are robust or subtle, *in vivo* within living cells. However, the earlier iteration of this technique, BIoID, suffered from prolonged labeling times and was limited to cytosolic labeling ^66^. In contrast, the miniTurbo-driven proximity labeling approach we employed facilitates the screening of potential weak interactors within the ER lumen ^28^. The integration of miniTurbo with mass spectrometry has enabled the identification of a significantly enriched pool of proteins involved in redox reactions and protein glycosylation within the ER, encompassing P4HB, TXNDC5, PDIA3, PRKCSH, GANAB, and UGGTI. Among the array of potentially intriguing interactors with STIMs unveiled by our proteomic analysis, our attention was drawn to PRKCSH and GANAB. Serving as the regulatory and catalytic subunits of glucosidase II, respectively, PRKCSH and GANAB emerged as focal points of investigation. Herein, we present compelling evidence implicating Glu II as the specific enzyme responsible for further modifying STIM1’s glycosylation site, as depicted in Figures 1, 2, and 3.

Through the ablation of N-linked glycosylation via N131C/N171C (2NC), N131D/N171Q (DQ), or N131Q/N171Q (2NQ) mutants with STIM1, prior investigations underscore the pivotal roles of N131 and N171 in regulating STIM1 activation^19,67^. Cumulative evidence suggests that N-glycosylated STIM1 exhibits heightened sensitivity to fluctuations in ER Ca^2+^ levels, rendering STIM1 ideally poised as a binary activator of SOCE to fulfill cytosolic Ca^2+^ demands. Nonetheless, these mutations completely eliminate the oligosaccharide side chains, providing solely a glimpse into the effects of either final versions of glycosylated STIM1 or the consequences of glycosylation deficiency on STIM1. Further clarification regarding the role of STIM1 post the initial addition of the glycan (Glc3Man9GlcNAc2), prior to the cleavage of the two innermost glucose residues by Glu II, remains elusive. In essence, the physiological significance of Glu II in STIM1 extends beyond its role in protein folding quality control.

Given the localization of glycosylation sites within the EF-SAM domain of STIM1, modifications to glycosylation by Glu II likely exert effects on its Ca^2+^ binding affinity. Leveraging our advanced FRET tools developed, we were able to investigate this phenomenon under *in situ* conditions, mirroring physiological settings. Initially, we examined the impact of STIM1 mutants (2NQ) that completely eliminate glycosylation. Our *in situ* measurements unveiled that complete deglycosylation enhances STIM1’s Ca^2+^ affinity (Fig. 4A and 4B). The heightened Ca^2+^ affinity induced by the 2NQ mutant prolongs the residence of more STIM1 proteins in a resting state under specific stimulus conditions, contrary to the active state expected for STIM1 WT proteins. Consequently, this leads to a reduction in the STIM1:Orai1 ratio, resulting in an inhibitory effect on SOCE. Our data offer compelling support for the inhibition of SOCE by 2NQ, as documented in previous studies^67^.

Knocking down either PRKCSH or GANAB markedly reduces the Ca^2+^ affinity of STIM1 WT (Fig. 4F and 4G). This finding implies that in the absence of cleavage of the two innermost glucose residues, glycosylated STIM1 exhibits diminished Ca^2+^ affinity, potentially rendering it more sensitive to fluctuations in ER Ca^2+^ levels. The cellular system has already evolved a STIM2 variant with lower Ca^2+^ affinity, poised to respond to subtle ER Ca^2+^ changes. Hence, we posit that the presence of Glu II serves to augment STIM1’s Ca^2+^ affinity by modifying its glycosylation, thereby delineating distinct functional territories for the two STIM proteins.

A noteworthy discovery reported last year reveals that PRKCSH, GANAB, and CLPTM1 (Cleft lip and palate transmembrane protein 1) form a ternary complex within the ER lumen, while GPER1 (The G protein-coupled estrogen receptor 1) directly interacts with CLPTM1 ^68^. Their findings also imply an indirect interaction between GPER1 and STIM1. However, they overlook the crucial relationship between PRKCSH, GANAB, and STIM1. Our data strongly suggest that this indirect interplay between STIM1 and GPER1 likely occurs via PRKCSH and GANAB.

The models depicted in Fig. 6 encapsulate several key findings of our study. Clearly, under physiological conditions, abundant Glu II catalyzes modifications at all N-glycosylation sites of STIM1. Upon store depletion, STIM1 proteins undergo activation, with Glu II further co-localizing with STIM1 to facilitate these modifications. This process substantially augments STIM1’s Ca^2+^ binding affinity, resulting in fewer STIM1 molecules remaining in an activated state. By bolstering STIM1’s Ca^2+^ binding affinity, Glu II orchestrates the regulation of SOCE to prevent Ca^2+^ overload (Fig. 5D and 5E). Conversely, under pathological conditions, exemplified by Glu II knockdown, insufficient Glu II levels result in incomplete modification of STIM1’s N-glycosylation sites. Consequently, these incompletely modified glycosylated STIM1 proteins exhibit reduced Ca^2+^ binding affinity, leading to heightened STIM1 activation in response to similar decreases in ER Ca^2+^ levels. Moreover, their enhanced gating ability to the Orai1 channel on the plasma membrane amplifies SOCE, as depicted in Fig. 5A and 5C. Our study underscores the intricate role of Glu II in fine-tuning STIM1’s response to ER Ca^2+^ fluctuations through targeted modifications at its N-glycosylation sites, thereby influencing its activation status.

In summary, our investigation illuminates the intricate interplay between Glu II (PRKCSH/GANAB) and STIM1, unveiling their potential significance in cellular calcium signaling. The MRH domain of PRKCSH and specific residue D524 within GANAB emerge as pivotal determinants governing their interaction with STIM1, underscoring their crucial roles in regulating calcium signaling dynamics. Our findings introduce a novel regulatory function for Glu II in modulating STIM1 activation dynamics, thereby influencing SOCE. These insights contribute to the expanding understanding of the molecular mechanisms orchestrating calcium signaling regulation. Comprehending the functional interplay between PRKCSH/GANAB and STIM1 offers valuable insights into the intricate network of proteins involved in calcium homeostasis and cellular signaling, with implications for both cellular physiology and pathology.

## Materials and Methods

### Plasmids construction

The cDNA clones encoding human GANAB (Ensembl CCDS8026.1, 944 aa), PRKCSH (Ensembl CCDS32911.1, 528 aa), and UGGT1 (Ensembl CCDS2154.1, 1555 aa) were synthesized meticulously by Qinglan Biotech. Correspondingly, the coding sequences (CDS) of mTurquoise2 and miniTurbo underwent precise synthesis by the same esteemed institution. Additionally, the mNeonGreen plasmid and a plasmid harboring the mScarlet sequence were generously provided by Dr. Chen Liangyi of Peking University. STIM11-310, amplified from wild-type STIM1^37^, and ECFP△C11 were seamlessly inserted into the pCDNA3.1 vector, which had been linearized by EcoRV and XhoI, utilizing a multiple-fragment homologous recombination kit (Cat#: C113, Vazyme Biotech, Nanjing, China). This process yielded the STIM11-310-ECFP△C11 construct. A comparable methodology was employed to produce mNG△N5-SOAR1L, PRKCSH-mScarlet, and GANAB-mScarlet constructs. miniTurbo, alongside either STIM1 or STIM2, was introduced into the pEYFP-N1 vector, linearized by EcoRI and BamHI, generating miniTurbo-STIM1-YFP or miniTurbo-STIM2-YFP constructs, through the same homologous recombination kit. Additionally, full-length wild-type PRKCSH (WT) and deletion mutants (△MRH, R139H, PRKCSH1-446, and PRKCSH1-413) tagged with a C-terminal mScarlet were integrated into the EcoRV and XhoI sites of the pcDNA 3.1 vector. Similar techniques were applied to produce various GANAB mutants. Verification of all plasmids was conducted through Sanger sequencing.

### Cell culture and transfection

HEK293 and HeLa cells (ATCC, catalog numbers CRL-1573 and CL-0101, respectively) were cultured routinely in DMEM (Cytiva) supplemented with 10% FBS (ExCell) and 1% penicillin-streptomycin at 37°C with 5% CO2 ^69^. Transfections of HEK293 cells were conducted via electroporation using the Bio-Rad Gene Pulser Xcell system (Bio-Rad, Hercules, CA, USA) in 4 mm cuvettes with OPTI-MEM medium. A voltage step pulse of 180 V for 25 ms was applied in 0.4 ml of medium, while for HeLa cells, the conditions were 260 V, 525 µF, in 0.5 ml of medium. N2a cell transfections involved purified plasmids and lipofectamine 3000 (Invitrogen, Waltham, MA, USA), following the manufacturer’s protocol. Transfected cells were seeded onto round coverslips, initially cultured in serum-free OPTI-MEM for 40 minutes (Thermo Scientific, Waltham, MA, USA), followed by regular DMEM supplemented with 10% FBS and 1% penicillin-streptomycin for 24 hours.

To generate stable cells, the Ca^2+^ indicator TurNm^53^ was transfected into HEK293 cells. Following selection with 100 μg/ml G418 for 5-7 days, cells were diluted to single clones and cultured further. Healthy clones showing robust expression and normal Ca^2+^ responses were chosen. HEK-miniTurbo-STIM1-YFP and Neuro-2a-miniTurbo-STIM2-YFP stable cells were established using identical procedures.

### Gene knockdown by shRNA

Knockdown of PRKCSH, GANAB, or UGGT1 gene expression in HeLa-SK cells (STIM1 and STIM2 double knockout) utilized lentiviral shRNA vectors. PRKCSH shRNA (5’-CGATGACTATTGCGACT GCAA-3’) was inserted into the pLKO.1 vector between AgeⅠ and EcoRⅠ sites. Similar methods generated GANAB shRNA (5’-CCCAACCTCTTTGTCTGGAAT-3’) and UGGT1 shRNA (5’-CCTGTTTACCTCTCTGGCTAT-3’). A nonspecific scrambled shRNA (5’-GAGGTAGTCTTAGAGGGTTGA-3’) in pLKO.1 served as control. Cells were transfected with shRNA via electroporation. Gene silencing was confirmed by qPCR and western blotting 48 hours post-transfection.

### Total RNA isolation and quantitative real-time polymerase chain reaction (qPCR) analysis

Total RNA extraction from HeLa-SK cells employed TRIzol, followed by reverse transcription using PrimeScript™ RT Master Mix (Takara, cat. no. RR036A) per manufacturer’s instructions. The resulting cDNA served as a template for quantitative PCR (qPCR), combined with primers (Table 1) and SYBR Green PCR Master Mix (GenStar Biosolutions, cat. no. A314). Quantitative PCR (qPCR) was conducted on a QuantStudio™ 6 Flex Real-Time PCR System (Applied Biosystems, CA, USA). Relative mRNA levels were determined using the Comparative Ct (△△CT) method and normalized to human GAPDH expression levels ^25^.

**Table 1.**
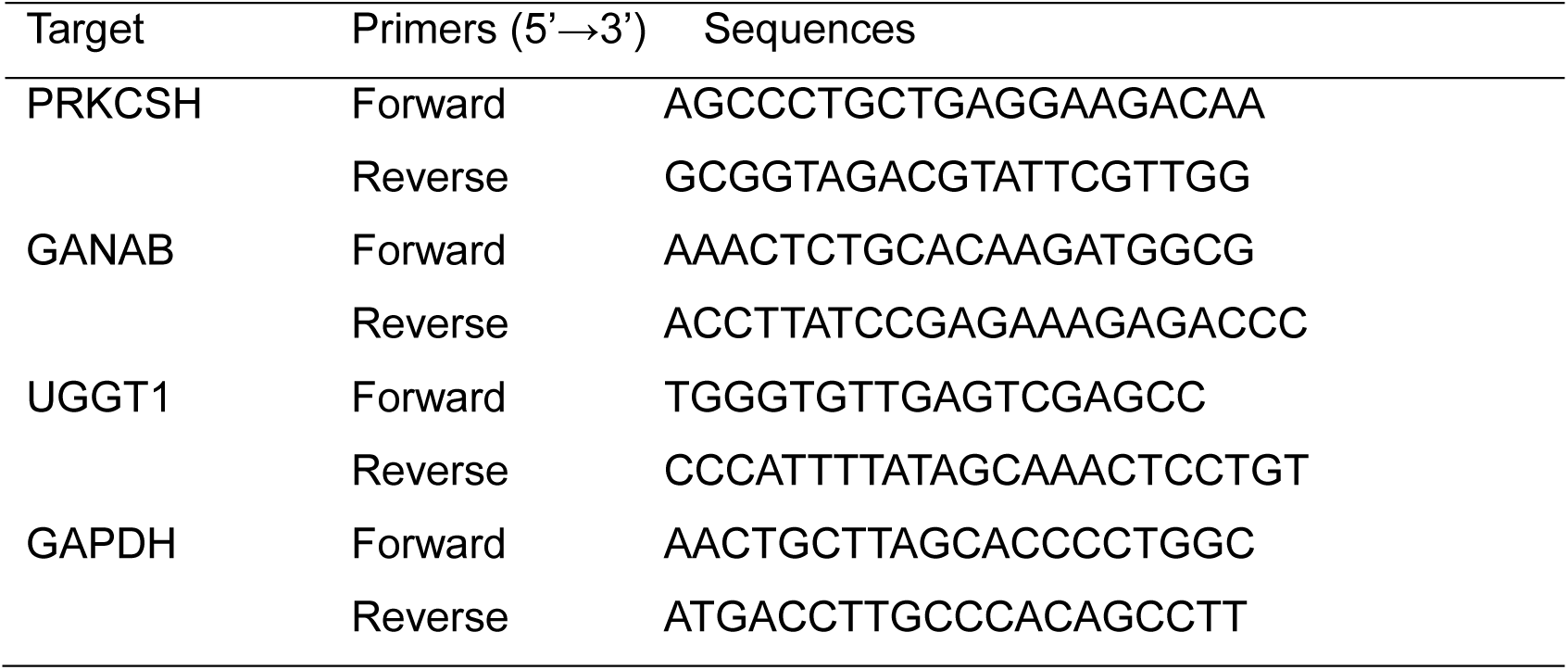
Sequences of primers used for qPCR.

### Single-cell intracellular Ca^2+^ imaging

All Ca^2+^ imaging experiments were executed following established protocols ^70,71^. Utilizing a ZEISS observer-Z1 microscope equipped with an X-Cite 120-Q light source (Lumen Dynamics, Waltham, MA, USA), ORCA-Flash4.0 V3 Digital CMOS camera (Hamamatsu, Japan), and 40× oil objective (NA=1.30) with standard Semrock filters, experiments were conducted at room temperature. Control of the imaging system was facilitated by SlideBook6.0.23 software (Intelligent Imaging Innovations, Inc.). The basic imaging solution comprised 107 mM NaCl, 7.2 mM KCl, 1.2 mM MgCl2, 11.5 mM glucose, and 20 mM Hepes-NaOH, pH 7.2, with additional reagents incorporated as indicated.

Intracellular Ca^2+^ imaging experiments primarily utilized HEK 293 cells, which were engineered to stably express TurNm, a recently introduced Ca^2+^ indicator ^53^. Imaging was conducted at a frame rate of one frame per 2 seconds using the TurNm filter set, comprising mTurquoise2 (mTq2): 438 ± 12 nm Ex, 483 ± 16 nm Em, and NEMOm: 500 ± 12 nm Ex, 542 ± 13 nm Em ^72^. Fluorescence data were exported and processed using Matlab 2023b (The MathWorks, Natick, MA, USA) to determine the relative ratio of FNEMOm/FmTurquoise2, indicating cytosolic Ca^2+^ concentration changes, followed by visualization using Prism 9.5.1. Traces depicted represent findings from a minimum of three independent experiments, each involving 30-60 individual cells.

### Förster Resonance Energy Transfer (FRET) imaging and measurements of Ca^2+^ affinity of STIM1

The FRET assay employed the same imaging system utilized in previous Ca^2+^ imaging experiments, with calibrations and offline analyses performed following established protocols ^73–75^. Two pairs of fluorescent proteins (FPs) were employed: CFP&YFP and ECFP△C11&mNG△N5. Images were captured every 10 seconds, capturing emissions from CFP/ECFP△C11 (438 ± 12 nm _Ex_/483 ± 16 nm _Ex_), YFP/mNG△N5 (510 ± 5 nm _Ex_ /542 ± 13.5 nm _Em_), and FRET raw (438 ± 12 nm _Ex_ /542 ± 13.5 nm _Em_). Parameters for the CFP-YFP FRET pair remained consistent with previous studies ^75^, while recalibration was conducted for the ECFP△C11&mNG△N5-mediated FRET. The recalibration and computation of ECFP△C11&mNG△N5-mediated FRET involved the formula: FRET_c_ = F_raw_ − F_d_/D_d_ × F_ECFP△C11_ − F_a_/D_a_ × F_mNG△N5_. Here, F_d_/D_d_ denotes the quantified bleed-through of ECFP△C11 into the FRET filter (0.73), while F_a_/D_a_ represents the quantified bleed-through of mNG△N5 through the FRET filter (0.36). Normalized FRET was computed by normalizing FRET_c_ against F_ECFP△C11_ to alleviate expression level discrepancies. Fluorescence data were extracted from SlideBook6.0.23 and analyzed using Matlab R2023b. Representative traces from a minimum of three independent experiments, each encompassing 15-30 cells, were presented as mean ± s.e.m. These meticulous methodologies ensure accurate assessment and interpretation of the FRET dynamics in the cellular context.

*In situ* or *in cellulo* Ca^2+^ titrations of STIM were performed using HeLa SK cells. For *in cellulo* measurements, cells transiently co-expressed YFP-SOAR1L with either PM-SC2222-CFP or PM-SC2222-N222Q-CFP. These were observed using Ca^2+^ imaging buffer solutions (as described above) containing 8 different free Ca^2+^ concentrations, ranging from zero to 30 mM Ca^2+^, to obtain response ratios through FRET imaging. Similarly, *in situ* measurements were conducted in cells transiently co-expressing mNG△N5-SOAR1L with either STIM11-310-ECFP△C11 or variants. These measurements utilized a solution containing 10 mM NaCl, 140 mM KCl, 1mM MgCl2, 20 mM HEPES, 0.025 mM digitonin, 0.01 mM ionomycin, and 1 mM EGTA (pH 7.4). During measurements, cells were permeabilized with the above solution containing different free Ca^2+^ concentrations, ranging from zero to 2 mM Ca^2+^, to obtain response ratios by FRET imaging. Ca^2+^ affinities of the STIM1 or variants were then calculated by fitting the FRET-Ca^2+^ relationship to the Hill equation using Prism 9.5.1 software. All experiments were carried out at room temperature. Traces shown are representative of at least three independent repeats, with 30-60 single cells analyzed per each repeat.

### Confocal microscopy

The ZEISS LSM880 system, equipped with a 63x oil objective (NA 1.4) and controlled by ZEN 2.1 software, was utilized for confocal imaging. CFP or mTurquoise2, YFP/mNeonGreen, and mScarlet were excited using 405 nm, 488 nm, and 543 nm lasers, respectively, and detected within the ranges of 420-500 nm, 470-540 nm, and 590-690 nm. The slice thickness was maintained at 1 μm. Image analysis was performed using Image J software (NIH). Each experiment was conducted a minimum of three independent transfections, with representative data presented.

### Western Blotting

Total proteins were extracted using the Total Protein Extraction Kit (BB18011; BestBio) following the manufacturer’s instructions, and their concentration was measured by the BCA protein assay kit (E162-01; GenStar). Six times loading buffer was added to the protein extracts, which were then boiled for 5 minutes at 98°C. Twenty micrograms of total protein were separated on 10% SDS-PAGE and transferred onto PVDF membranes (Millipore). The resulting membranes were blocked for 1 hour at 37°C with TBST buffer (12 mM Tris-HCl, pH 7.5, 137 mM NaCl, 2.68 mM KCl, 0.1% Tween 20) containing 5% nonfat dried milk or bovine serum albumin (BSA), then incubated with primary antibody overnight at 4°C. After washing three times (10 minutes each) in TBST buffer, the membranes were loaded with secondary antibody for 40 minutes at room temperature. Detection was performed by ECL (GS009-4; Millipore) solution and imaged using the Tanon5200 detection system finally. HRP-labeled Streptavidin, STIM1, PRKCSH, GANAB, and UGGT1 were detected with anti-HRP-labeled Streptavidin (N100, Thermo Fisher scientific, USA), anti-STIM1 antibody (5668S; CST; 1:1000 dilution), anti-PRKCSH antibody (Proteintech, Cat:12148-1-AP; 1:1000), anti-GANAB antibody (Proteintech, Cat:29183-1-AP; 1:1000), and anti-UGGT1 antibody (Proteintech, Cat:14170-1-AP; 1:1000) followed by anti-rabbit-IgG (7074S; CST; 1:4000 dilution) respectively. Internal control β-actin was detected with anti-β-actin antibody (CW0096; CWBIO: 1:4000 dilution) and corresponding secondary antibody is anti-mouse-IgG (7076S; CST; 1:5000 dilution). The representative data shown were carried out from three independent experiments. The intensity of the images was quantified by Image J software, and the resulting data were plotted with prism9.5.1 software.

### Biotinylation, enrichment of biotinylated proteins and silver staining

Biotin (Sangon Biotech; 58-85-5) was solubilized in DMSO, sterilized via filtration, and stored at -20°C in 100- to 250-µL aliquots. Cells were exposed to 50 µM biotin for 3 hours, unless otherwise stated. After incubation, the media was removed, and cells were thrice rinsed with cold PBS. Subsequently, cells were harvested, and total proteins were extracted using the Total Protein Extraction Kit (BB18011; BestBio). The protein samples underwent dialysis to eliminate free biotin, followed by determination of protein concentration using a BCA protein assay (E162-01; GenStar). Streptavidin agarose beads (11206D; Invitrogen) were utilized for enriching biotinylated proteins, with 100 µl of a 50% slurry used per sample. After washing and equilibration, protein samples were added to the beads and rotated for 1.5 hours for enrichment. The beads were washed five times with 1× PBST buffer containing protease inhibitors. Following this, the samples were suspended in 2 × SDS sample buffer, and the proteins were eluted by boiling for 5 minutes at 100°C. The separated biotinylated proteins were resolved using 10% Bis-Tris protein gels. SDS-PAGE was conducted, and the gels were subjected to Silver Stain Kit (LC6070; Thermo Fisher scientific) staining according to the provided instructions. A parallel SDS-PAGE was employed for Western blotting to detect biotinylated proteins using streptavidin-HRP (1:4000). Subsequently, the protein lanes were excised and submitted to Beijing HuaDa protein company for mass spectrometry identification, resulting in the elucidation of the proteome associated with STIM.

### Statistical analysis

All quantitative data are presented as means ± SEM derived from a minimum of three independent biological replicates. Comparisons between two groups were scrutinized using unpaired t-tests, while comparisons among multiple groups underwent analysis through One-way ANOVA.

## Acknowledgements

This work was supported by the Ministry of Science and Technology of China (2019YFA0802104 to Y.W.), the National Natural Science Foundation of China (91954205 and 92254301 to Y.W.).

## Author contributions

Y.W. and Y.Z. supervised and coordinated the study. Y.W. and Y.Z. designed the experiments. Y.D. designed and generated all the plasmid constructs. Y.D. performed most live cell Ca^2+^ imaging, FRET imaging and confocal experiments. S.Z. performed a few Ca^2+^ imaging experiments. F.W. performed a few FRET imaging experiments. J.L. and X.Y. prepared the proteomic samples and performed the mass spectrometry analyses. Y.D. performed all the other experiments and did data analysis. Y.Z., Y.W. and Y.D. wrote and revised the manuscript with inputs from all the other authors.

## Conflicts of interests

All authors declare no conflicts of interests.

## Supplementary Figures

**Figure S1.**
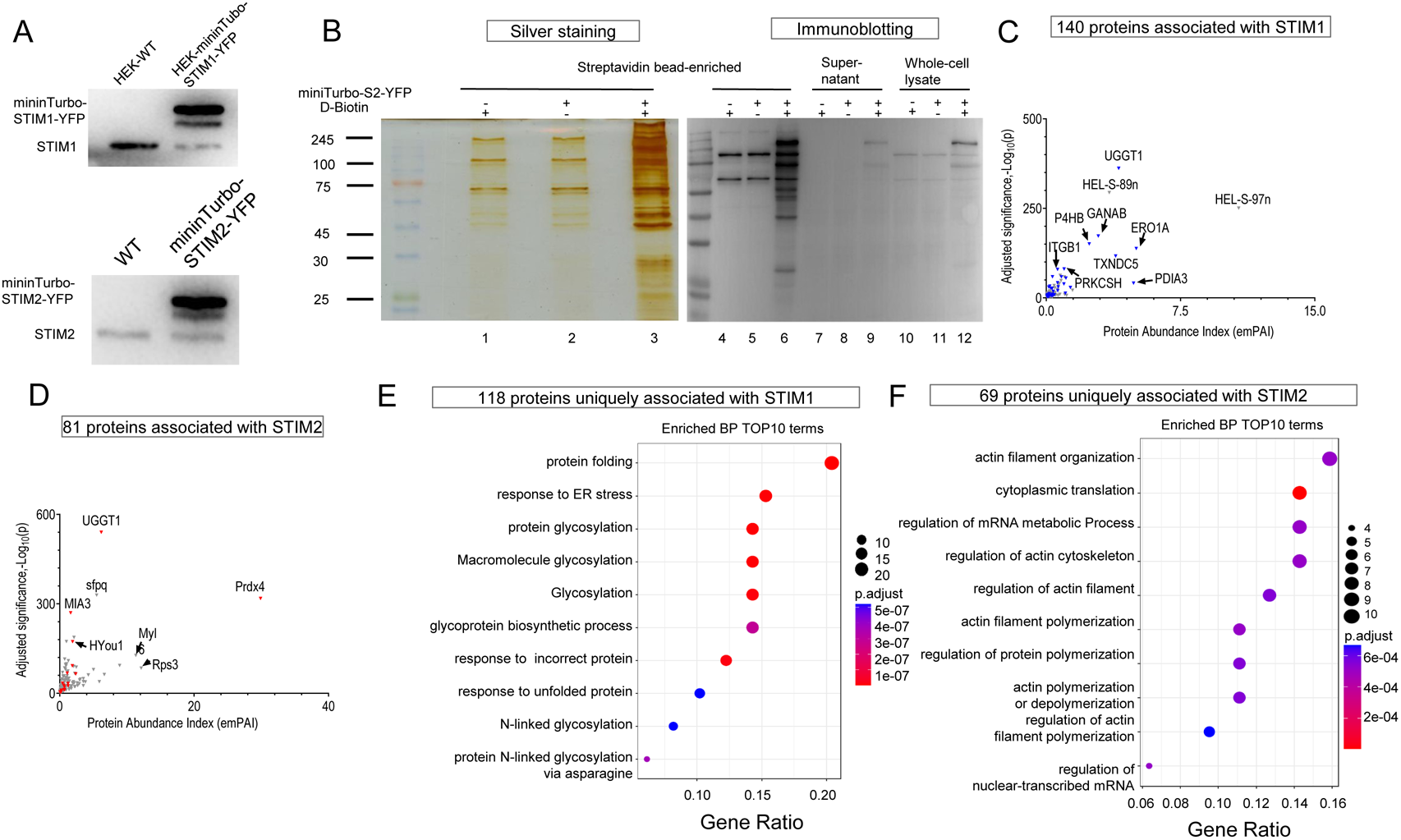
Proteomic mapping, bioinformatics analysis and dynamic co-localization of STIM-proximal proteomics. (A) Western blot analysis confirmed the expression of STIM1 and miniTurbo-STIM1-YFP (top), as well as STIM2 and miniTurbo-STIM2-YFP (bottom). (B) Silver staining (left) or immunoblotting (right) revealed biotinylated proteins adjacent to STIM2. Enriched samples are displayed on SDS-PAGE (lanes 1–3), while immunoblots detected with streptavidin-HRP show enriched samples (lanes 4–6), supernatant samples (lanes 7–9), and whole cell lysate (lanes 10–12). (C-D) Volcano plots illustrate the quantified proteome adjacent to STIM1 (C) or STIM2 (D), with colored dots indicating ER-localized proteins. (E-F) Bubble charts depict the top 10 enriched biological processes (BP) of the proteome adjacent to STIM1 or STIM2 identified in (C) or (D), respectively.

**Figure S2.**
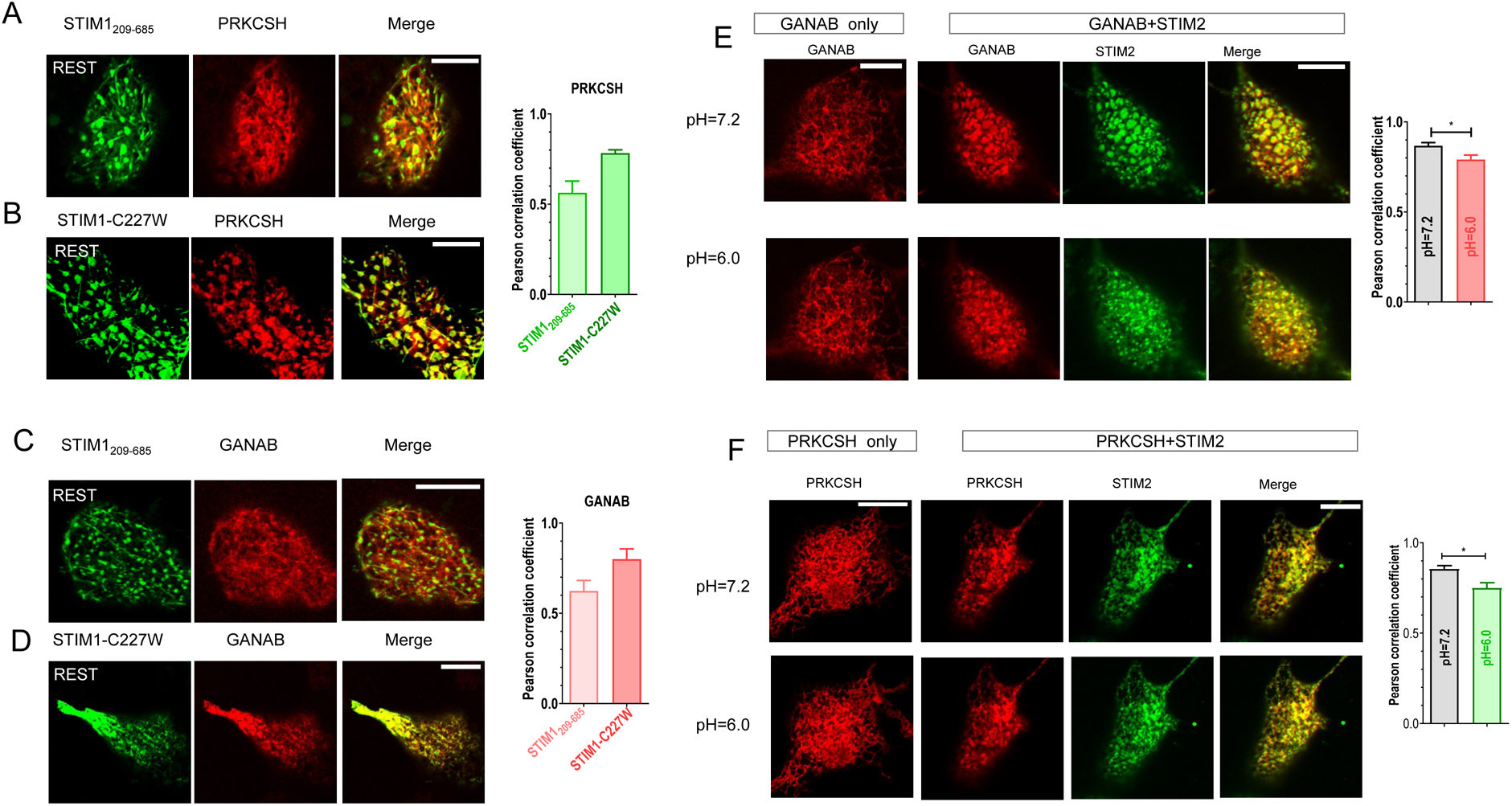
Super-resolution confocal imaging of HEK293 cells co-expressing STIM1_209-685_ or mutated mTurquoise2-STIM1, PRKCSH or GANAB. Images of cells under resting conditions and following a 5-minute depletion of ER-Ca^2+^ stores using 2.5 µM IONO were captured from the identical field of view. Presented are typical cellular images from three independent experiments, with more than 15 cells examined each time. In the images, STIM1 variants are depicted in green, while PRKCSH or GANAB variants are represented in red. (A-B) Illustrate colocalization between STIM1_209_-_685_ (A) or STIM1-C227W (B) and PRKCSH. Bar chart on the right shows statistical analysis indicating the extent of co-localization between PRKCSH and STIM1_209--685_ or STIM1-C227W. C-D) Depict colocalization between STIM1_209-685_ (C) or STIM1-C227W (D) and GANAB, complemented by a bar chart offering statistical analysis of their co-localization. (E-F) Confocal-imaging assay results demonstrate GANAB’s dynamic co-localization with STIM2 protein upon intracellular acidification in HEK293 cells, complemented by a bar chart offering statistical analysis of their co-localization. Scale bar: 10 µm.

**Figure S3.**
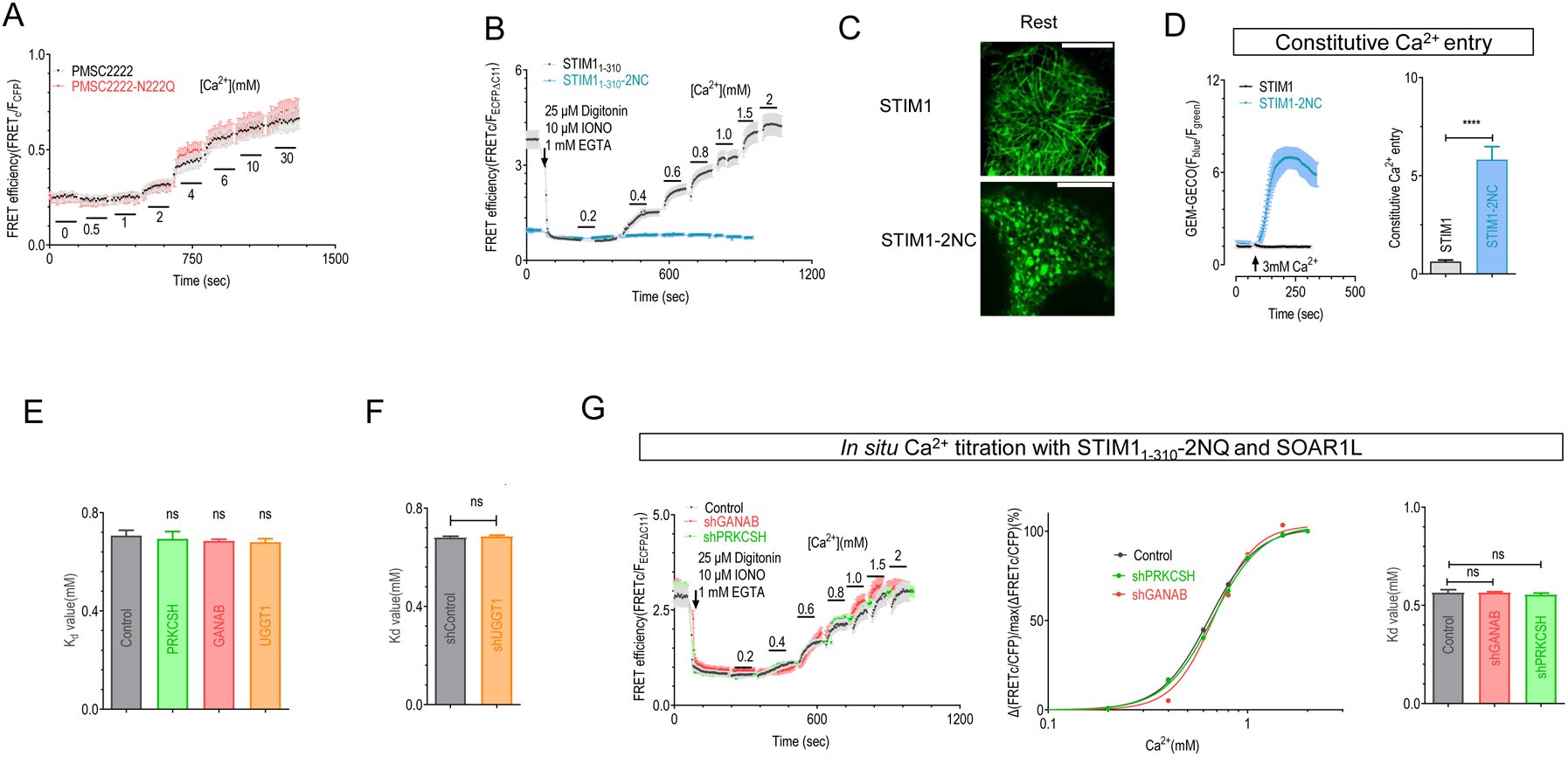
The effect of expression changes of GANAB, PRKCSH or UGGT1 on STIM1’s Ca^2+^ affinity. (A) Intracellular Ca^2+^ titration traces were depicted through FRET signals between co-expressed YFP-SOAR1L and PM-SC2222-CFP/PM-SC2222-N222Q-CFP. B) Ca^2+^-dependent FRET responses were observed in HeLa-SK cells transiently co-expressing either mNGΔN5-SOAR1L with STIM1_1-310_-ECFPΔC11 or with STIM1_1-310_-N131C-ECFPΔC11. C) Representative images illustrating the distribution of mTurquoise2-STIM1 or mTurquoise2-STIM1-2NC in cells. D) Cytosolic Ca^2+^ response. Left panel displays typical traces; right panel presents statistical analysis. E) Statistical analysis of the effects of PKRCSH, GANAB, or UGGT1 co-expression on the Kd value of STIM1 in Ca^2+^ binding (n = 3, unpaired Student’s t-test, ns, not significant, with more than 30 cells examined in each group). F) Statistical analysis of the Kd value of STIM1 in Ca^2+^ binding with or without UGGT1 knockdown (KD) (n = 3, unpaired Student’s t-test, ns, not significant, with more than 30 cells examined per measurement). G) *In situ* Ca^2+^ titration curves regarding FRET responses between mNG△N5-SOAR1L and STIM1_1-310_-2NQ-CFP△C11 constructs in cells transfected with control shRNA, shPRKCSH, or shGANAB. The left panel illustrates traces of Ca^2+^ response; the middle panel presents Ca^2+^-titration curves; the right panel displays statistical analysis (n = 3, unpaired Student’s t-test, ns, not significant, with more than 30 cells examined per measurement).

**Figure S4.**
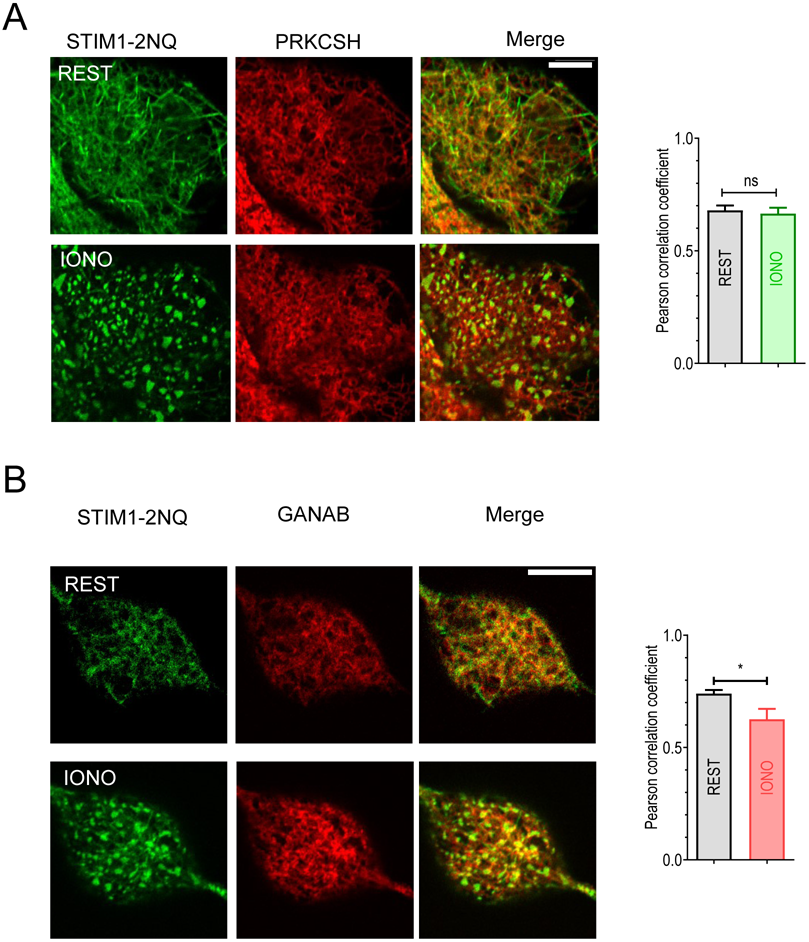
STIM1-2NQ abolishes its dynamic co-localization with GANAB or PRKCSH. Cellular imaging was conducted under basal conditions and after a 5-minute depletion of ER-Ca^2+^ stores using 2.5 µM IONO, capturing images from the same field of view. Presented are typical cellular images from three independent experiments, each involving examination of more than 15 cells. STIM1-2NQ is visualized in green, while PRKCSH or GANAB is depicted in red. The scale bar denotes 10 µm. (A) showcases the co-localization between STIM1-2NQ and PRKCSH, along with a corresponding bar chart providing statistical insights into their co-localization extent. (B) The co-localization between STIM1-2NQ and GANAB, complemented by a bar chart offering statistical analysis of their co-localization.

**Figure S5.**
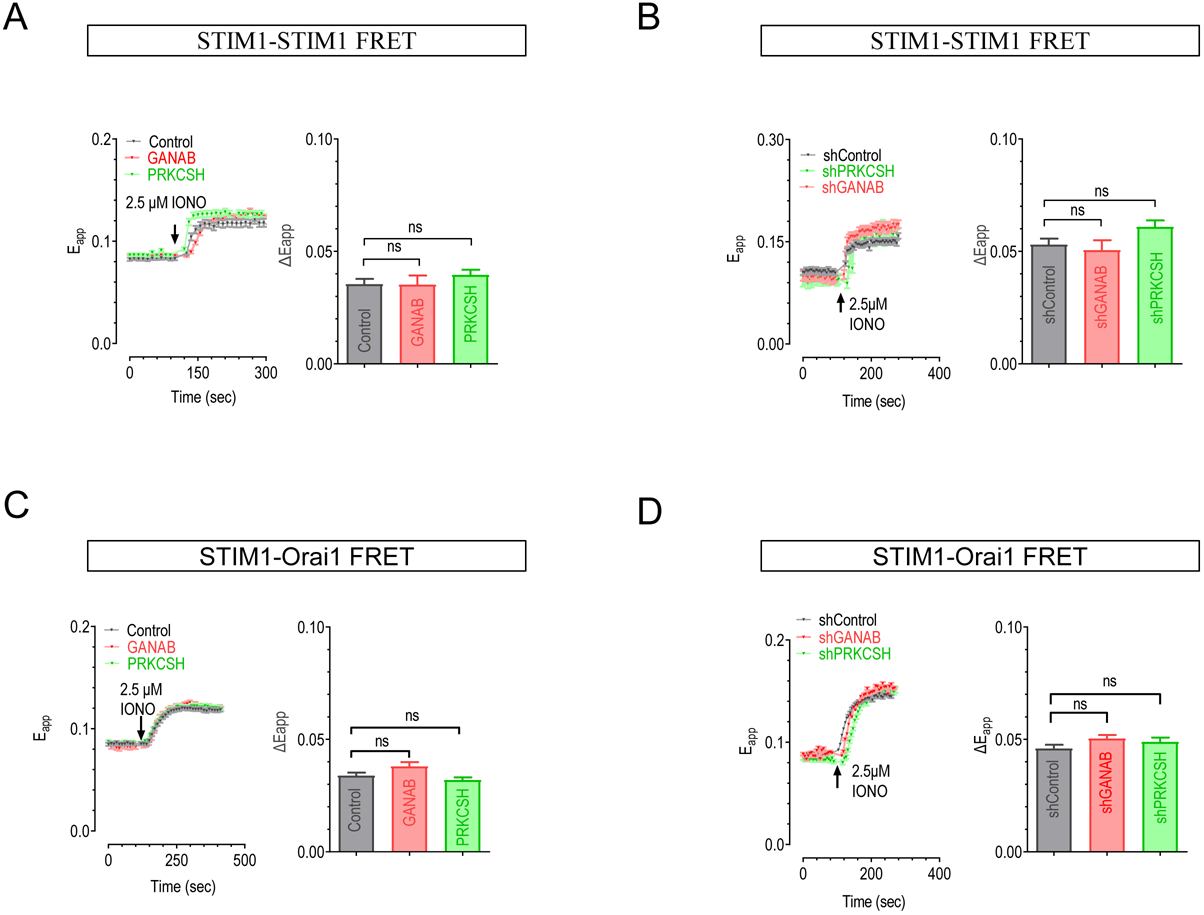
Absence of Influence from GANAB, PRKCSH, or UGGT1 on STIM1-STIM1 and STIM1-Orai1 Interactions. (A-D) Administration of 2.5 µM ionomycin induced ER Ca^2+^ store depletion. The left side illustrates the time course, while the right side displays the quantification of △Eapp. △Eapp values were derived by subtracting the resting Eapp from the plateau Eapp observed 5 minutes post-ionomycin treatment. (A-B) Impact of PRKCSH or GANAB overexpression or knockdown on FRET signals between co-expressed CFP-STIM1 and YFP-STIM1 in cells. (C-D) Influence of PRKCSH or GANAB overexpression or knockdown on FRET signals between STIM1-CFP and Orai1-YFP co-expressed in cells (n = 3, with over 25 cells per measurement, assessed via unpaired Student’s t-test, ns, indicating no significance).

## Notes

### Competing Interest Statement

The authors have declared no competing interest.

### Summary of Updates

This version changes the order of authors, with Yandong Zhou listed as the corresponding author

